# Single-Cell Imaging Maps Inflammatory Cell Subsets to Pulmonary Arterial Hypertension Vasculopathy

**DOI:** 10.1101/2022.12.03.518033

**Authors:** Selena Ferrian, Aiqin Cao, Erin F. Mccaffrey, Toshie Saito, Noah F. Greenwald, Mark R. Nicolls, Trevor Bruce, Roham T. Zamanian, Patricia Del Rosario, Marlene Rabinovitch, Michael Angelo

**Affiliations:** Department of Pathology, Stanford University, Stanford CA, 94305, USA; Department of Pediatrics, Division of Cardiology, Stanford University, Stanford CA, 94305, USA; Division of Pulmonary, Allergy, and Critical Care Medicine and; Vera Moulton Wall Center for Pulmonary Vascular Disease, Stanford CA, 94305, USA; Veterans Affairs Palo Alto Health Care System, Palo Alto CA, 94305, USA; Early Clinical Development, ECDi, Genentech Inc., South San Francisco CA, 94080, USA

**Keywords:** Pulmonary arterial hypertension, vascular remodeling, MIBI-TOF, monocyte-derived dendritic cells (mo-DCs), indoleamine 2, 3-dioxygenase 1 (IDO-1), TIM-3+ T cells, neutrophils, interferon gamma (IFN-γ), SAM and HD domain-containing deoxynucleoside triphosphate triphosphohydrolase 1 (SAMHD1), bone morphogenetic protein receptor 2 (*BMPR2*) mutation

## Abstract

**Rationale:** Elucidating the immune landscape within and surrounding pulmonary arteries (PAs) is critical in understanding immune-driven vascular pathology in pulmonary arterial hypertension (PAH). Although more severe vascular pathology is often observed in hereditary (H)PAH patients with *BMPR2* mutations, the involvement of specific immune cell subsets remains unclear.

**Methods:** We used cutting-edge multiplexed ion beam imaging by time-of-flight (MIBI-TOF) to compare PAs and adjacent tissue in PAH lungs (idiopathic (I)PAH and HPAH) with unused donor lungs.

**Measurements:** We quantified immune cells’ proximity and abundance, focusing on those linked to vascular pathology, and evaluated their impact on pulmonary arterial smooth muscle cells (SMCs) and endothelial cells (ECs).

**Results:** Distinct immune infiltration patterns emerged between PAH subtypes, with intramural involvement independently linked to PA occlusive changes. Notably, we identified monocyte-derived dendritic cells (mo-DCs) within PA subendothelial and adventitial regions, influencing vascular remodeling by promoting SMC proliferation and suppressing endothelial gene expression across PAH subtypes. In HPAH patients, pronounced immune dysregulation encircled PA walls, characterized by heightened perivascular inflammation involving TIM-3+ T cells. This correlated with an expanded DC subset expressing IDO-1, TIM-3, and SAMHD1, alongside increased neutrophils, SMCs, and α-SMA+ECs, reinforcing the severity of pulmonary vascular lesions.

**Conclusions:** This study presents the first architectural map of PAH lungs, connecting immune subsets not only with specific PA lesions but also with heightened severity in HPAH compared to IPAH. Our findings emphasize the therapeutic potential of targeting mo-DCs, neutrophils, cellular interactions, and immune responses to alleviate severe vascular pathology in IPAH and HPAH.

## INTRODUCTION

Pulmonary arterial hypertension (PAH), a progressive disease, is classified into etiology-based subtypes. Hereditary (H)PAH often arises from mutations in bone morphogenetic protein receptor type 2 (*BMPR2*), while idiopathic (I)PAH lacks an identifiable cause. Despite varying underlying pathologies among subtypes, current FDA-approved treatments offer limited improvement, resulting in nearly 40% mortality over 5 years (1).

Chronic inflammation and altered immunity have been increasingly recognized in PAH pathogenesis (2,3). Monocyte and macrophage recruitment (4,5), T- and B cell infiltration (2), tertiary lymphoid structures near pulmonary arteries (PAs) (9), regulatory T cell loss and dysfunction (6,7) along with increased DC subset activation (8, 10, 11) and neutrophil involvement (10) in PA and lung tissue, have been associated with PAH initiation and worsening.

Despite emerging immunotherapies targeting individual cell types or compounds (12), the multifaceted nature of PAH involves numerous pathways and cellular processes. Our study comprehensively maps the immune cell landscape in PAs, exploring microenvironmental complexities, vascular cell interactions, and their contribution to disease onset and progression. We hypothesize that specific immune cell phenotypes may predict the extent of vascular damage and differentiate *BMPR2* loss-of-function mutations in HPAH, linked to severe vascular pathology (13), heightened inflammation (14,15), and increased mortality (16) from IPAH. This insight could shed light on cellular mechanisms driving disease progression and guide the development of tailored therapies.

High throughput technologies, such as multiplexed ion beam imaging by time-of-flight (MIBI-TOF) (17), facilitate the study of multiple cells, their activation, and spatial arrangement within tissues. This enhances our understanding of cell interactions during PA remodeling, advancing PAH understanding, and potentially guiding innovative multi-targeted treatments. Using MIBI-TOF, our study images a panel of thirty-five proteins, encompassing immune cell differentiation and function, as well as tissue-specific markers. We present the first architectural map of PAH lung tissue, linking immune cell subsets to PA lesions and highlighting their role in the severe pathology of HPAH. Monocyte-derived DCs (mo-DCs) located within the PA subendothelial and adventitial regions drive vascular remodeling and endothelial modulation, independent of subtype. HPAH exhibits increased perivascular inflammation, marked by TIM-3+ T cells and neutrophils, further exacerbating the pathology. Targeting immune dysregulation and perivascular inflammation, including the mo-DCs, through targeted modulation of cellular interactions and immune responses, offers a promising strategy to alleviate progression in both PAH subtypes, with particular significance for HPAH.

Some of the results of these studies have been previously reported in the form of an abstract(s)(51–54).

## METHODS

An Expanded Methods section is available in the Online Data Supplement.

### Study Cohort

Lung tissues were obtained at transplant from PAH (n=12) and donor controls (Con, n=10) through the Pulmonary Hypertension Breakthrough Initiative and the Cardiovascular Medical Research and Education Fund. Peripheral blood mononuclear cells (PBMCs) were isolated from PAH and Con blood collected from the Stanford Biobank and Blood Bank, respectively.

### Histopathological Scoring

Approximately five distal pulmonary arteries (PAs) were selected per patient based on scoring of vascular morphology. See Figure 2A and E1B in the Online Data Supplement.

**Figure 1.**
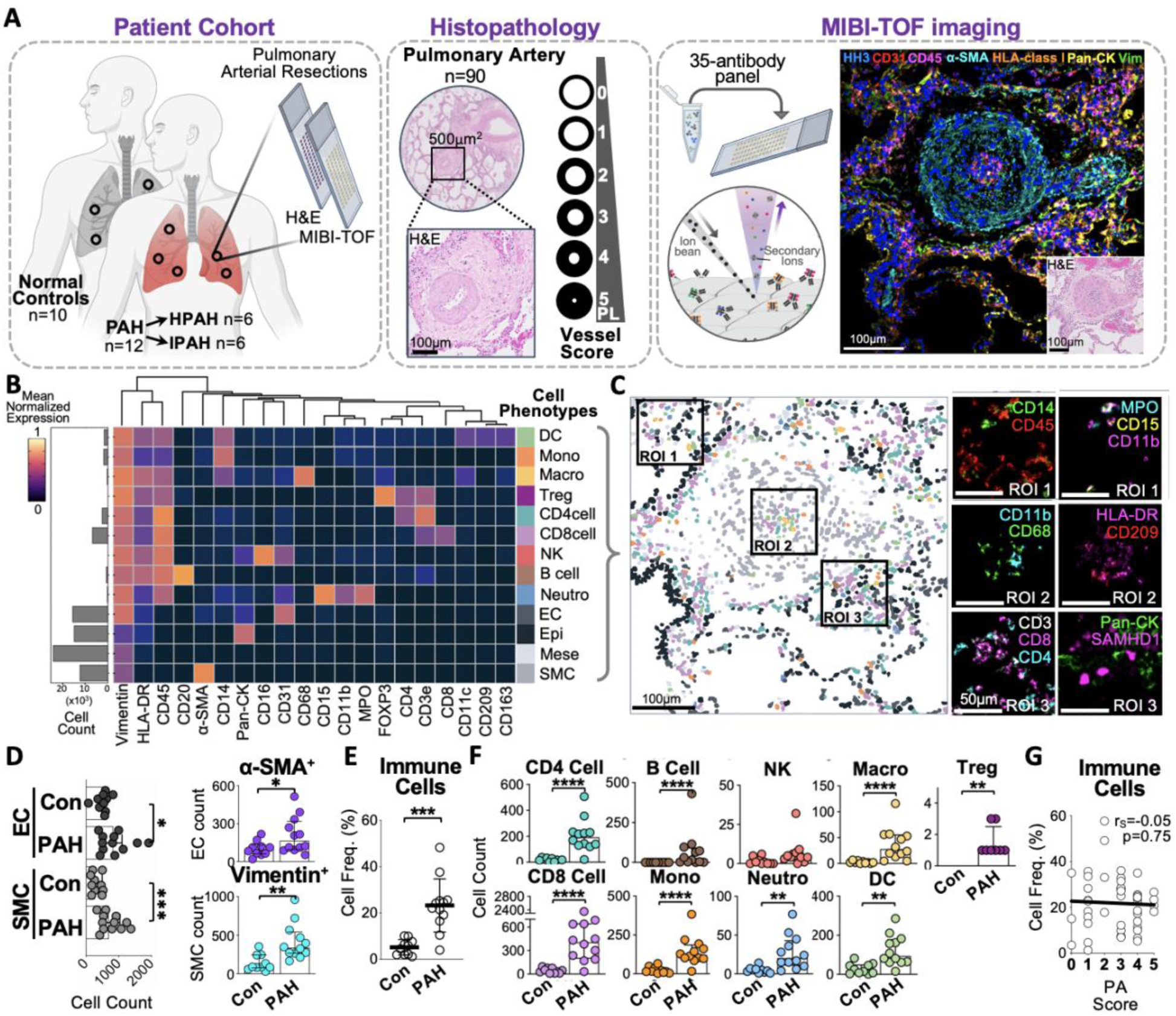
Multiplexed imaging interrogation of pulmonary arteries from pulmonary arterial hypertension patients and donor controls. (**A**) Schematic depicting the workflow for the study. Representative pulmonary arteries (PAs) from ten donor control (Con) and twelve pulmonary arterial hypertension (PAH) patients were selected and embedded in a tissue microarray, of which six patients were diagnosed with idiopathic (I)PAH and six with hereditary (H)PAH. Ninety PAs were scored on hematoxylin and eosin (H&E) in relation to the severity of the lesion and subsequent tissue sections were obtained to conduct MIBI-TOF analyses. PL, Plexogenic Lesion. The image was created using BioRender. (**B**) Cell lineage assignments based on mean normalized expression of lineage markers (heatmap columns) hierarchically clustered (Euclidean distance, average linkage). Rows are ordered by cell lineage (bottom) and immune cell breakdown (top). The absolute abundance of each cell type is displayed (left). (**C**) Representative MIBI-TOF image overlay from a PA section showing cell identity by color, as defined in *B*. Scale bars = 100μm. Six zoomed insets showing MIBI-TOF overlays across diverse regions displaying neutrophil cells (ROI 1), DCs and macrophages (ROI 2), T cells along with epithelial and SAMHD1+ cells (ROI 3). Scale bars = 50μm. (**D**) Difference in endothelial cells (EC) and smooth muscle cells (SMC) count between donor controls (Con) and pulmonary arterial hypertension (PAH) patients, along with difference in α-SMA+ECs and Vim+SMCs. Vim, Vimentin. (**E**) Difference in immune cells frequency (mean and SD) between donor controls (Con) and pulmonary arterial hypertension (PAH) patients. (**F**) Difference in immune cell subsets frequency (median and IQR) between donor controls (Con) and pulmonary arterial hypertension (PAH) patients. (**G**) Relationship (correlation) between immune cell infiltration and pulmonary artery (PA) score in pulmonary arterial hypertension (PAH) patients. Spearman’s rank correlation (rS) was employed for the correlation, with a 95% confidence interval and two-tailed test. α-SMA, alpha-smooth muscle actin; MPO, myeloperoxidase; HLA-DR, human leukocyte antigen – DR isotype; HH3, Histon H3; Pan-CK, Pan-cytokeratin; Vim, Vimentin; ROI, region of interest; EC, endothelial cell; Epi, epithelial cell; Mese, mesenchymal cell; SMC, smooth muscle cell and fibroblast; DC, dendritic cell; Mono, monocyte; Macro, macrophage; Neutro, neutrophil; Treg, regulatory CD3+CD4+FOXP3+ cell; NK, natural killer (NK) cell. P-values were calculated with unpaired t test with Welch’s correction or a Wilcoxon rank-sum test, where *p < 0.05, **p < 0.01, ***p < 0.001, ****p < 0.0001.

**Figure 2.**
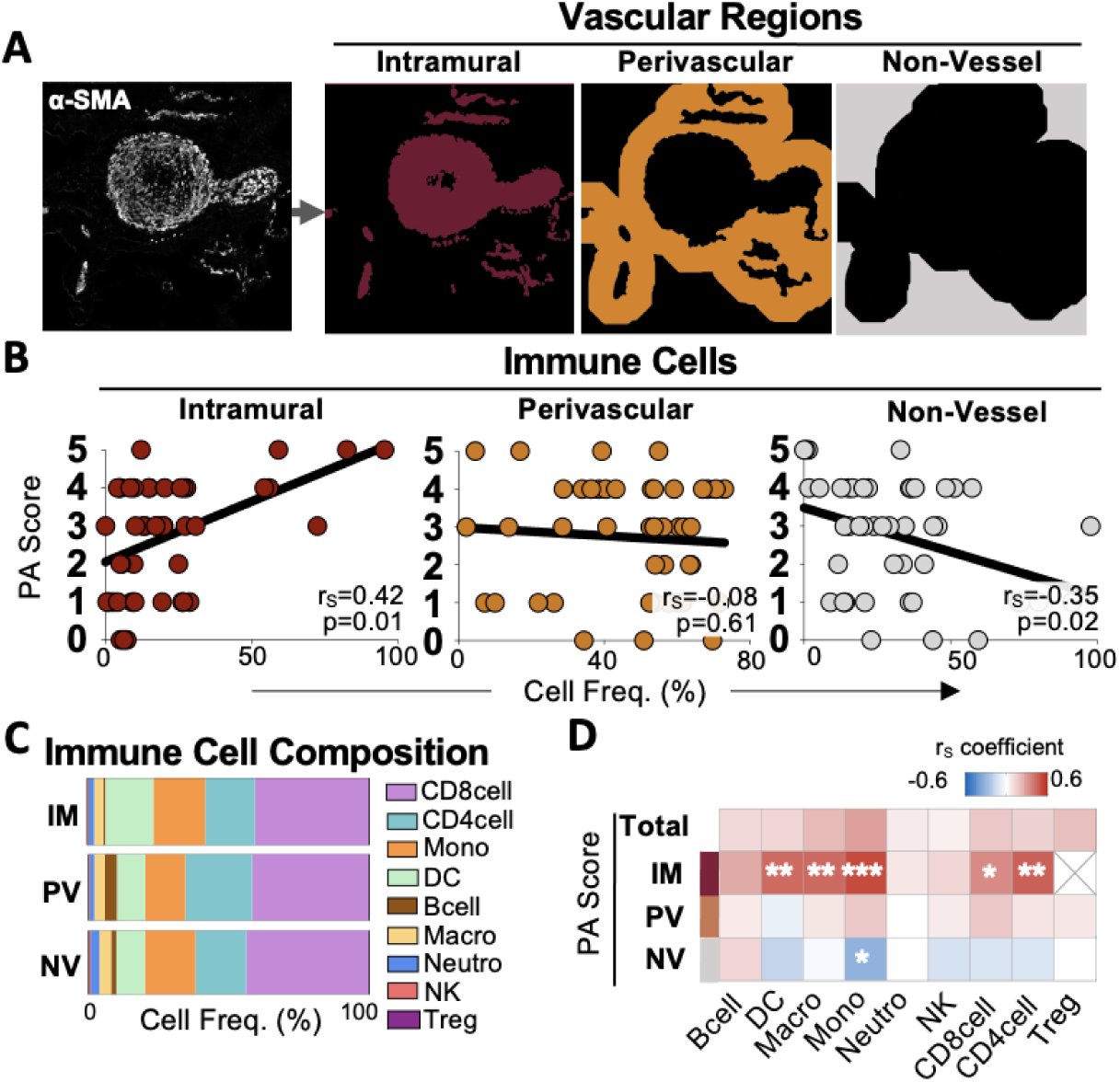
Intramural immune cells relate to exacerbated vascular pathology. (**A**) Binary masks were generated to localize immune cells with respect to the vascular architecture. α-SMA staining from the MIBI-TOF scan was used to extrapolate the intramural region (maroon). A region within 100 px (∼50 μm) of the vessel was defined as a ‘perivascular’ (dark orange). Any region beyond that was considered ‘non-vessel’ (grey). (**B**) The relationship (correlation) between pulmonary artery (PA) score and the immune cell frequency across vascular regions in patients with pulmonary arterial hypertension (PAH). (**C**) The immune cell composition across vascular regions, with ‘IM’ representing intramural, ‘PV’ representing perivascular, and ‘NV’ representing non-vessel regions. (**D**) Heatmap displaying the relationship (correlation) between immune cell counts across vascular regions (heatmap rows) and pulmonary artery (PA) score (heatmap columns). The heatmap includes total cell counts (regardless of vascular region) as well as counts specific to intramural (IM), perivascular (PV), and non-vessel (NV) regions. The p-values are indicated by white asterisks. Correlations were carried out using Spearman’s rank correlation (rS) with 95% confidence interval and two-tailed test, where *p < 0.05, **p < 0.01, ***p < 0.001, ****p < 0.0001. α-SMA, alpha-smooth muscle actin; HH3, histone H3; DC, dendritic cell; Mono, monocyte; Macro, macrophage; Neutro, neutrophil; Treg, regulatory CD3+CD4+FOXP3+ cell, NK, natural killer (NK) cell.

### Reagents

See Table E1 in the Online Data Supplement.

### Tissue staining and imaging

The tissue was stained with a 35-marker panel and imaged using MIBI-TOF as previously described (17). See Figure 1A and E1C, F in the Online Data Supplement.

### Image Processing and Segmentation

Image processing and segmentation followed established methods (17). See Figure E1D-E in the Online Data Supplement.

### Cell Profiling and Functional Analysis

Single-cell data were processed as described (18) and analyzed for cell phenotypes using the Bioconductor "FlowSOM" package (19). Functional protein positivity thresholds were determined using the "MetaCyto" package (20). See Figure 1B-C.

### Vascular Region Masking

Three vascular regions were defined: intramural (IM) with α-SMA staining, perivascular (PV) within 100 pixels of the vessel edge, and beyond that as non-vessel (NV). See Figure 2A in the Online Data Supplement.

### Spatial Analyses

Cell-to-vessel distances were calculated, and tissue microenvironments were created by analyzing cell neighborhoods within a 25 μm radius of index cells. See Figure 3A and E3A in the Online Data Supplement.

**Figure 3.**
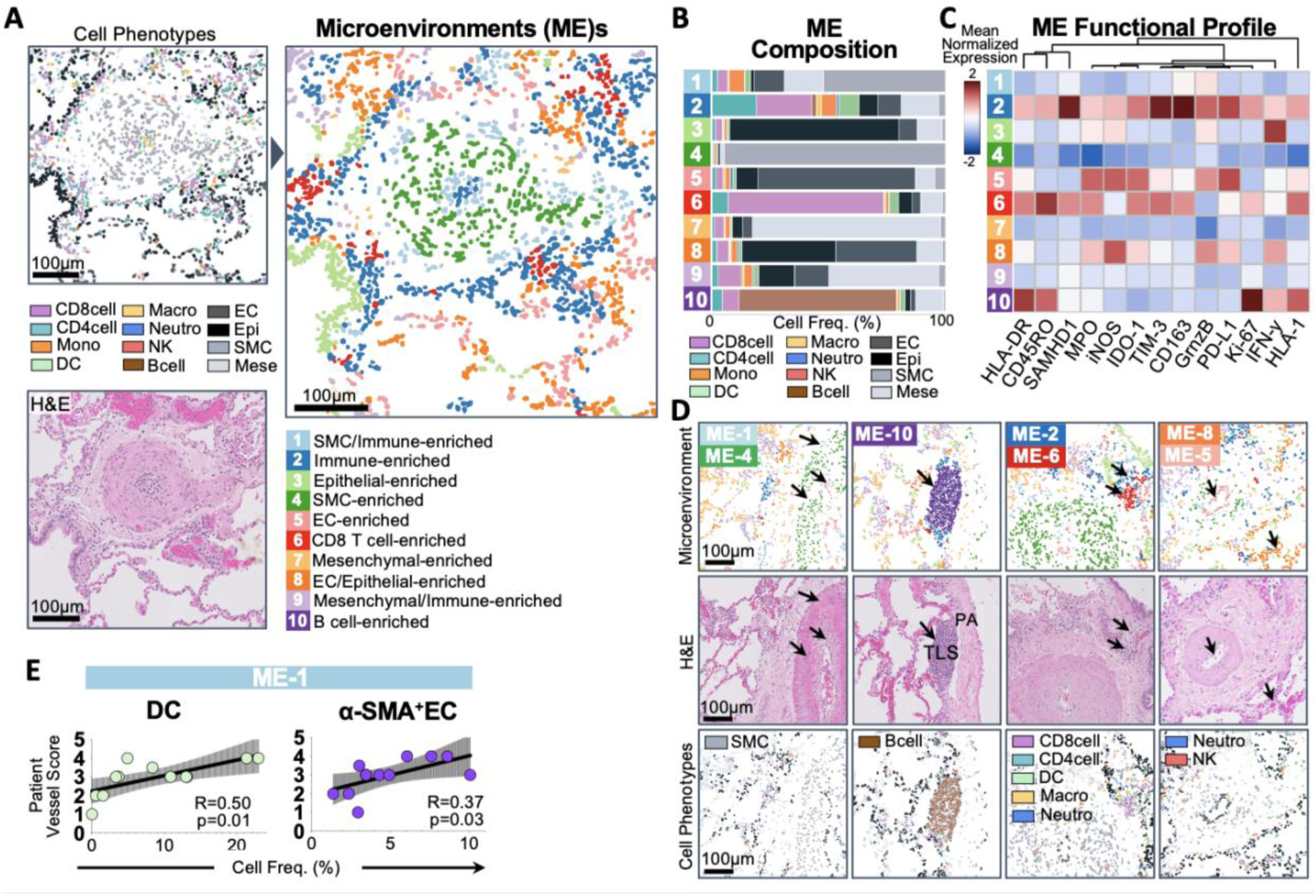
The spatial location of DCs and *α*-SMA+ endothelial cells relate to vessel score. (**A**) Representative image of a pulmonary artery (PA) displaying how cell types are grouped into local microenvironments (MEs). The corresponding hematoxylin and eosin (H&E) image, provides context for the PA tree architecture. Ten microenvironments are displayed in the image corresponding, to different combinations of proximal cell types, as defined in *B*. Scale bars = 100μm. (**B**) Cell composition of the ten tissue microenvironments (MEs) identified: immune and smooth muscle cell-enriched (ME-1); immune-enriched (ME-2); epithelial-associated (ME-3); SMC-enriched (ME-4); endothelial-enriched (ME-5); CD8 T cell-enriched (ME-6); mesenchymal-associated (ME-7); endothelial and epithelial-enriched (ME-8); immune and mesenchymal-enriched (ME-9); and B cell-enriched (ME-10) microenvironments. (**C**) Heatmap showing the functional profile of each microenvironment (ME). The mean normalized expression of hierarchically clustered (Euclidean distance, average linkage) functional markers across microenvironments (MEs) is reported. GrnzB, Granzyme B; MPO, Myeloperoxidase; HLA-I, human leukocyte antigen class I; HLA-DR, human leukocyte antigen – DR isotype. (**D**) Representative images of pulmonary arteries (PAs) showing their architecture with hematoxylin and eosin (H&E) staining and the cell composition of specific microenvironments (MEs). Black arrows highlight the ME locations within the images. Both ME-1 and ME-4 are intramural, with ME-1 in the subendothelial and adventitial regions, and ME-4 in the medial regions of the PA. ME-10, predominantly B cell-enriched, is adjacent to the PA wall and localizes in tertiary lymphoid structures (TLSs). ME-2 and ME-6, enriched with immune cells, surround the PA wall. Both ME-5 and ME-8 are localized within endothelial-enriched regions, with ME-5 in the intima layer and ME-8 in extramural vessels surrounding the PA, enriched with neutrophils and NK cells. Scale bars = 100μm. (**E**) Relationship (linear regression) between the frequency of dendritic cells (DCs) or α-SMA+ endothelial cells (EC) within ME-1 and the patient vessel score in pulmonary arterial hypertension (PAH). The p-values are reported as follows: *p < 0.05, **p < 0.01, ***p < 0.001, ****p < 0.0001. DC, dendritic cell; Mono, monocyte; Macro, macrophage; Neutro, neutrophil; NK, natural killer (NK) cell.

### Pulmonary Arterial SMC and EC growth

PASMCs were harvested from explanted lungs of Con patients and cultured in SMC growth medium, while ECs (PromoCell) were cultured in EC medium.

### Generation and Co-Culture of Monocyte Derived-Dendritic Cells

CD14+ cells from a Con individual and an IPAH patient were isolated from PBMCs using CD14 Microbeads, differentiated into mo-DCs (CellXVivo Human Monocyte-derived DC Differentiation Kit), and then co-cultured with PASMCs in 5% FBC SMC medium or ECs in EC medium for 72 hours.

### Pulmonary Arterial SMC Proliferation

After co-culturing, PASMCs proliferation was assessed using the CellTiter 96® AQueous One Solution Cell Proliferation Assay (MTS assay).

### Exosomes Isolation and Co-Culture

Exosomes were isolated (by ultracentrifuge) from plasma of three Con and six PAH patients, purified using CD66b/CEACAM 8-conjugated Dynabeads and quantified (Fluorocet Exosome Quantitation kit). Endothelial cells were cultured in EC medium and treated with an optimal dose of CD66b+ or CD66b-exosomes (10,000 count/cell), followed by a 72-hour incubation period before assessing gene expression.

### Real-time PCR Assessed mRNA Expression

SMC and EC total RNA were extracted (Qiazol), followed by reverse transcription and quantification using the 2^−ΔΔCt method, normalized to B2M.

### Enzyme-Linked Immunosorbent Assay (ELISA)

Signaling molecules were quantified in cultured media by ELISA following the manufacturer’s recommendations.

### Statistical Analysis

The Shapiro-Wild test gauged data normality. For two-group comparisons, we employed Mann-Whitney or two-tailed Welch’s T-tests. Before evaluating co-culture differences, data was normalized to PASMC or EC control. Co-culture significance was assessed via a mixed-effects model, accounting for repeated measures as a random effect, followed by pairwise comparisons. Spearman’s rank correlation and ordinal logistic regression appraised predictor-vessel score associations.

## RESULTS

### Mapping Vascular Lesions in Pulmonary Arterial Hypertension

To understand the immune landscape in PAH pathology, we analyzed tissue sections from explanted lungs of PAH patients (n=12) and unused donor lungs (Con, n=10) obtained during transplantation (**Figure 1A**). The cohort included six IPAH and six HPAH patients, with summarized demographic and clinical profiles detailed in **Table 1**. We examined multiple regions of interest (ROIs) containing distal pulmonary arteries (PAs) characterized by lesions of varying severity (**Figure E1A-B** in the Online Data Supplement) to provide a more comprehensive understanding of the disease process.

**Table 1.**
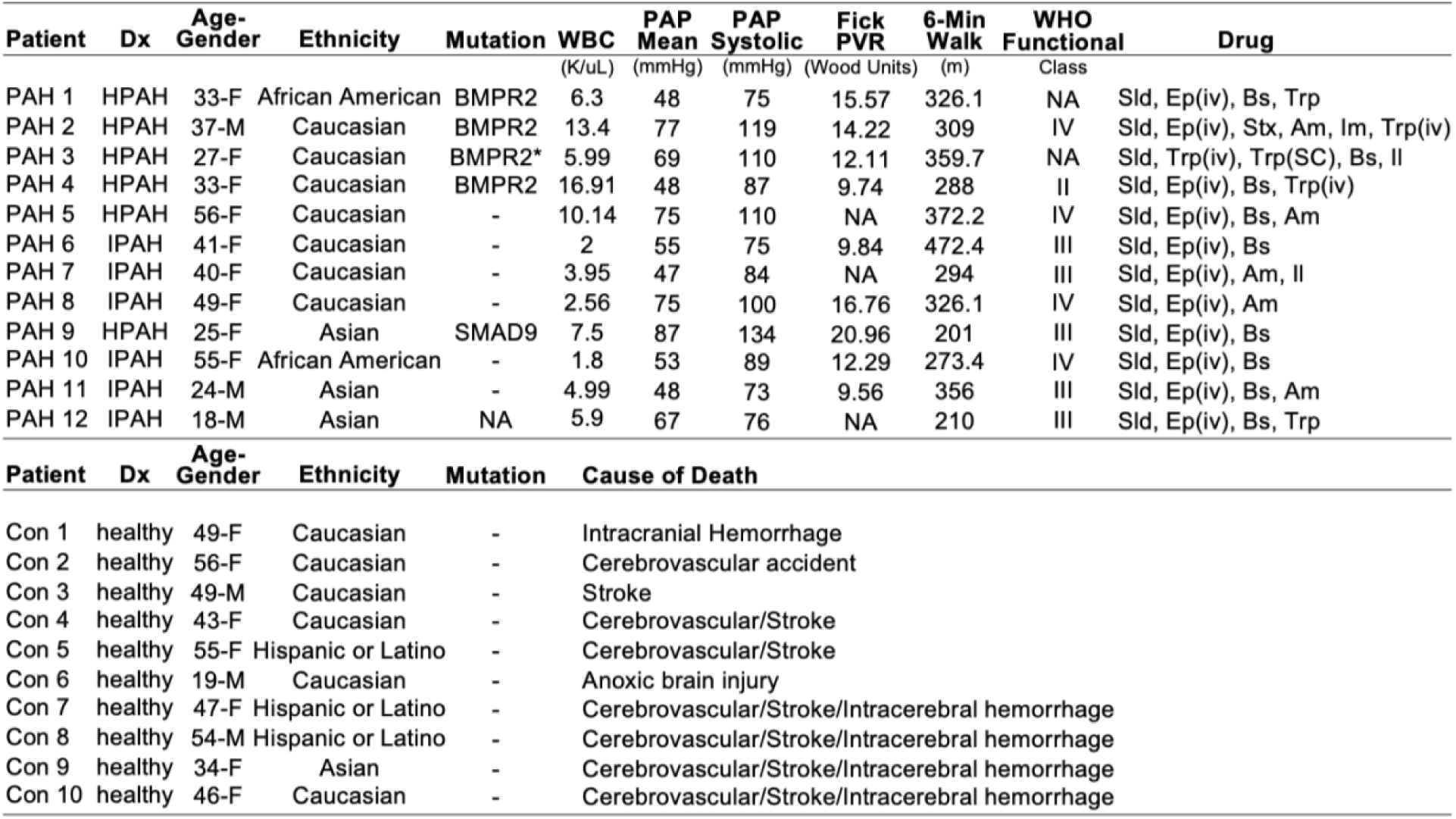
Clinical and demographic characteristics of patients. Clinical and demographic data were obtained from catheterization study performed closest to transplantation. Dx; Diagnosis: Pulmonary arterial hypertension (PAH) classification: (I)PAH, Idiopathic PAH; (H)PAH, hereditary PAH, known mutation as stated. Patient 5 (PAH 5) did not have a known mutation associated with PAH but had a significant family history of the condition. PAP, pulmonary arterial pressure. PVR, Pulmonary vascular resistance. Data were obtained from catheterization study performed closest to transplantation. 6 Min Walk = distance walked in six minutes. Data were obtained from study performed closest to transplantation. WHO Functional Class Functional based on the symptoms when doing everyday activities (I-IV, IV severe). Data were obtained from study performed closest to transplantation. PAH medications (Drug): Medications are listed according to total drug exposure during treatment period up to transplantation, not necessarily in combination. *Variant of unknown significance. Drugs abbreviations: Ep (Epoprostenol), Bs (Bosentan), Sld (Sildenafil), Trp (Treprostinil), Stx (Sitaxsentan), Am (Ambrisentan), Im (Imatinib), Il (Iloprost). iv (intravenous).

TABLE LEGEND
| Patient | Dx | Age-Gender | Ethnicity | Mutation | WBC<br>(K/uL) | PAP Mean<br>(mmHg) | PAP Systolic<br>(mmHg) | Fick PVR<br>(Wood Units) | 6-Min Walk<br>(m) | WHO Functional<br>Class | Drug |
| --- | --- | --- | --- | --- | --- | --- | --- | --- | --- | --- | --- |
| PAH 1 | HPAH | 33-F | African American | BMPR2 | 6.3 | 48 | 75 | 15.57 | 326.1 | NA | Sld, Ep(iv), Bs, Trp |
| PAH 2 | HPAH | 37-M | Caucasian | BMPR2 | 13.4 | 77 | 119 | 14.22 | 309 | IV | Sld, Ep(iv), Stx, Am, Im, Trp(iv) |
| PAH 3 | HPAH | 27-F | Caucasian | BMPR2* | 5.99 | 69 | 110 | 12.11 | 359.7 | NA | Sld, Trp(iv), Trp(SC), Bs, II |
| PAH 4 | HPAH | 33-F | Caucasian | BMPR2 | 16.91 | 48 | 87 | 9.74 | 288 | II | Sld, Ep(iv), Bs, Trp(iv) |
| PAH 5 | HPAH | 56-F | Caucasian | - | 10.14 | 75 | 110 | NA | 372.2 | IV | Sld, Ep(iv), Bs, Am |
| PAH 6 | IPAH | 41-F | Caucasian | - | 2 | 55 | 75 | 9.84 | 472.4 | III | Sld, Ep(iv), Bs |
| PAH 7 | IPAH | 40-F | Caucasian | - | 3.95 | 47 | 84 | NA | 294 | III | Sld, Ep(iv), Am, II |
| PAH 8 | IPAH | 49-F | Caucasian | - | 2.56 | 75 | 100 | 16.76 | 326.1 | IV | Sld, Ep(iv), Am |
| PAH 9 | HPAH | 25-F | Asian | SMAD9 | 7.5 | 87 | 134 | 20.96 | 201 | III | Sld, Ep(iv), Bs |
| PAH 10 | IPAH | 55-F | African American | - | 1.8 | 53 | 89 | 12.29 | 273.4 | IV | Sld, Ep(iv), Bs |
| PAH 11 | IPAH | 24-M | Asian | - | 4.99 | 48 | 73 | 9.56 | 356 | III | Sld, Ep(iv), Bs, Am |
| PAH 12 | IPAH | 18-M | Asian | NA | 5.9 | 67 | 76 | NA | 210 | III | Sld, Ep(iv), Bs, Trp |

**Table 1. Clinical and demographic characteristics of patients.**
| Patient | Dx | Age-Gender | Ethnicity | Mutation | Cause of Death |
| --- | --- | --- | --- | --- | --- |
| Con 1 | healthy | 49-F | Caucasian | - | Intracranial Hemorrhage |
| Con 2 | healthy | 56-F | Caucasian | - | Cerebrovascular accident |
| Con 3 | healthy | 49-M | Caucasian | - | Stroke |
| Con 4 | healthy | 43-F | Caucasian | - | Cerebrovascular/Stroke |
| Con 5 | healthy | 55-F | Hispanic or Latino | - | Cerebrovascular/Stroke |
| Con 6 | healthy | 19-M | Caucasian | - | Anoxic brain injury |
| Con 7 | healthy | 47-F | Hispanic or Latino | - | Cerebrovascular/Stroke/Intracerebral hemorrhage |
| Con 8 | healthy | 54-M | Hispanic or Latino | - | Cerebrovascular/Stroke/Intracerebral hemorrhage |
| Con 9 | healthy | 34-F | Asian | - | Cerebrovascular/Stroke/Intracerebral hemorrhage |
| Con 10 | healthy | 46-F | Caucasian | - | Cerebrovascular/Stroke/Intracerebral hemorrhage |

Leveraging MIBI-TOF imaging, we assessed thirty-five markers encompassing immune cell differentiation, activation, and tissue-specific characteristics. This approach aimed to elucidate intricate connections between immune cells and vascular pathology (**Figure 1A**; **Figure E1C** and **Table E1** in the Online Data Supplement), with detailed image processing and antibody validation outlined in **Figure E1D-F** in the Online Data Supplement.

Spatial analyses unveiled the interplay among immune, vascular, and non-vascular cells associated with vascular remodeling, including immune cell distribution, proximity to the PA wall, and intercellular distances (**Figure E1G** in the Online Data Supplement). MIBI-TOF enabled the identification and quantification of diverse cell phenotypes (**Figure 1B-C**).

Our analysis revealed a significant increase in the number and activation state of SMCs and ECs in PAH. Furthermore, the presence of α-SMA+ECs and Vimentin+SMCs pointed to a mesenchymal shift (**Figure 1D** and **E1H** in the Online Data Supplement). These findings, along with the observed increase in activated immune cell infiltration in PAH (**Figure 1E** and **E1H** in the Online Data Supplement), suggest immune-mediated interactions contributing to PAH progression. Regulatory T cells (Tregs) were sparse in the PAH cohort, while other immune cell types, except natural killer (NK) cells, exhibited an increase in PAH (**Figure 1F**). Further exploration identified tertiary lymphoid structures (TLSs) as partial contributors to the observed B cell increase (**Figure E1I** in the Online Data Supplement).

Notably, no correlation emerged between the extent of immune infiltration and the vascular pathology (**Figure 1G**). Moreover, circulating white blood cells (WBC) in PAH patients did not positively or negatively correlate with immune cell infiltration within the tissue (**Figure E1J** in the Online Data Supplement), suggesting a balance between cellular production, local recruitment, and potential amplification within the tissue—achieved through proliferation or prolonged half-life mechanisms.

### Intramural Immune Infiltration Correlates with Vascular Damage

We next categorized the immune infiltrate’s presence into three regions: intramural (IM), perivascular (PV), and nonvascular (NV) and assessed the regional relationship to the severity of PA lesions (**Figure 2A**). Immune cell counts were elevated across all PA regions in PAH compared to control PAs, with the intramural space exhibiting the most pronounced relative increase (**Figure E2A** in the Online Data Supplement). Notably, although immune cells were predominantly localized in the perivascular region (**Figure E2B** in the Online Data Supplement), only intramural immune cells positively correlated with PA score (**Figure 2B**). This correlation was especially notable for specific immune cell types, including DCs, monocytes, macrophages, and T cells, despite consistent immune cell composition across PA regions (**Figure 2C-D**). Further validation addressing within-patient scoring variability, demonstrated a robust association between intramural DCs and vessel score (**Figure E2C-D** in the Online Data Supplement).

In conclusion, our study underscores the significant association between intramural immune cells, especially DCs, and PA occlusion, highlighting the critical role of the microenvironment in DC function and its impact on PAH vascular pathology.

### Dendritic-Vascular Cell Interaction in the Progression of Vascular Remodeling

To gain insights into the individual or combined contribution of immune cells to vascular pathology, we conducted a ‘cell neighborhood analysis’ (**Figure E3A** in the Online Data Supplement). This approach allowed us to explore cell interactions in the PA microenvironment and create visualizations of the tissue, grouping cell types into local microenvironments (MEs) based on their phenotypic similarities and their proximity, within 25µm, to the index cells, as described in the Methods (**Figure 3A**). Ten distinct MEs were identified within the pulmonary tissue, each distinguished by varying compositions of cell types and functional profiles (**Figure 3B-D**). Notably, a tight interaction among immune cell types surfaced within ME-2 and ME-6, nestled in the immediate periphery of the PA wall. These microenvironments, abundant in T cells and DCs, displayed an activated and proliferative milieu (**Figure 3B-C** and **E3B** in the Online Data Supplement). SMCs held prominence within ME-4 (media) and displayed moderate presence within ME-1 (adventitia and subendothelium), alongside immune cells and ECs (**Figure 3B, 3D**). B cells formed TLSs in ME-10, adjacent to PAs, showcasing both proliferative prowess and antigen presenting capabilities (**Figure 3C-D** and **E3B** in the Online Data Supplement). Neutrophils and occasional NK cells localized preferentially in immune (ME-2) and endothelial (ME-5 and ME-8) enriched areas. ME-1, a subendothelial and adventitial microenvironment, emerged as a predictor of the PA score (**Table E2** in the Online Data Supplement). The presence of DCs and α-SMA+ECs within ME-1 strongly correlated with the degree of vascular pathology (**Figure 3E** and **E3C-D** in the Online Data Supplement). This finding suggests that the interaction between DCs and vascular cells may contribute to the progression of vascular remodeling.

### Mo-DCs: Orchestrators of Vascular Remodeling and Endothelial Dysfunction

To explore the DC profile associated with our findings, we identified four DC subsets (**Figure E4A** in the Online Data Supplement). Among them, CD11c-DC3s emerged as the predominant subset within the ME-1, displaying markers characteristic of monocyte-derived DCs (**Figure 4A-B** and **E4B** in the Online Data Supplement). Interestingly, these CD11c-DC3 cells correlated with vessel score and proliferative (Ki-67+) SMCs (**Figure 4C**) yet exhibited no correlation with hemodynamic factors or mesenchymal transition in SMCs and ECs (**Figure E4C-D** in the Online Data Supplement).

**Figure 4.**
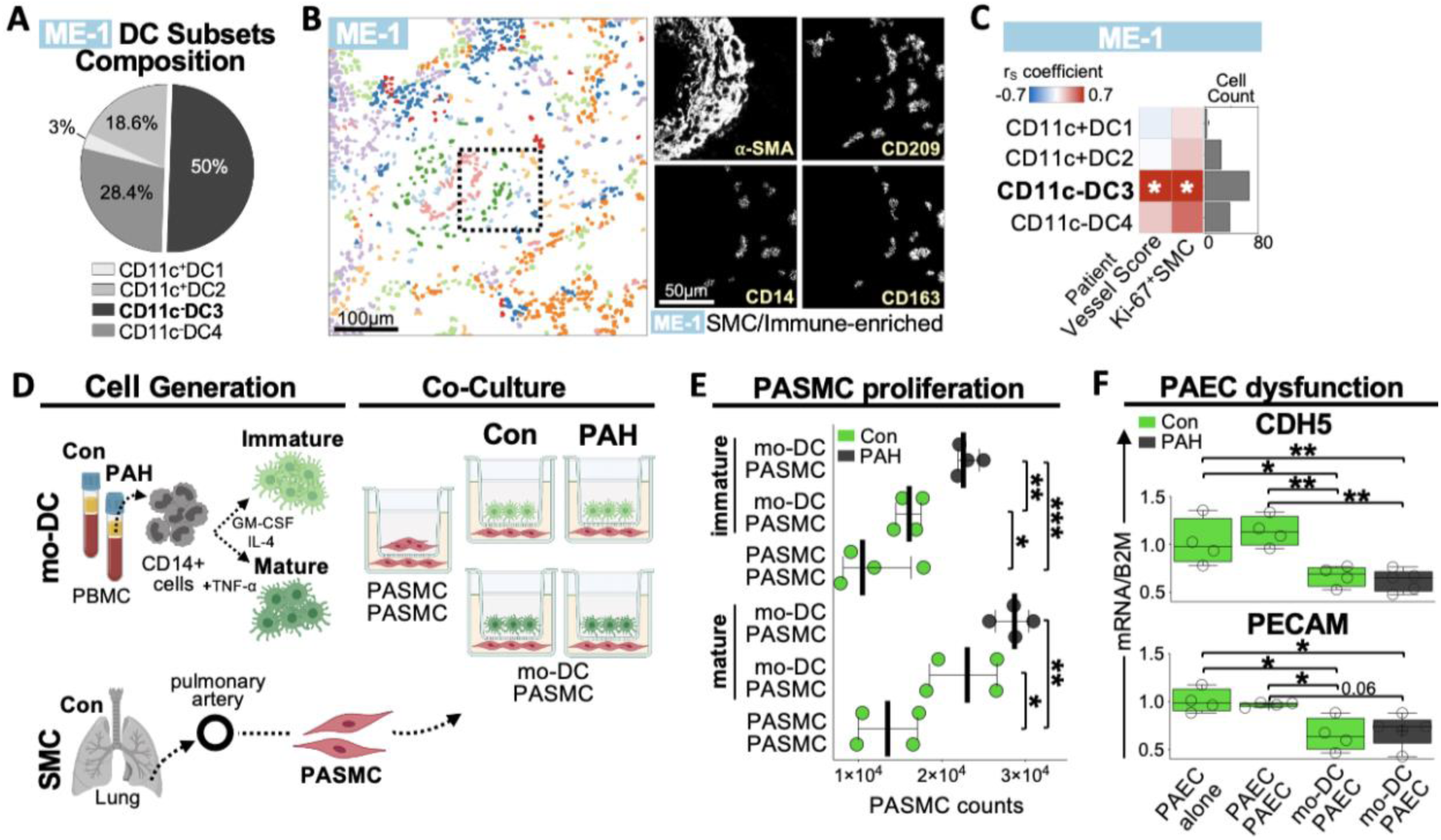
The spatial location of monocyte-derived dendritic cells dictates their involvement in vascular remodeling and endothelial dysfunction. (**A**) Dendritic cell (DC) composition within the subendothelial and adventitial regions (ME-1) of pulmonary arteries. Highlighted in bold are CD11c-DC3s (mo-DC-like cells), the predominant DC subset. (**B**) Representative image of a pulmonary artery showing the spatial localization of distinct microenvironments (MEs), denoted by color as in Fig. 3A. A zoomed inset of a region including ME-1 was chosen to examine the presence of CD11c-DC3s using CD14, CD209 and CD163 marker expression. α-SMA was displayed to highlight the pulmonary arterial tissue. ME map scale bar = 100μm; zoomed insets scale bars = 50μm. (**C**) Heatmap displaying the association (correlation) between dendritic cell (DC) subsets within ME-1 and patient vessel score or proliferating smooth muscle cells (Ki-67+SMC). The p-values (white asterix) are reported. The absolute abundance of each cell type is displayed (right). Correlations were carried out using Spearman’s rank correlation (rS) with 95% confidence interval and two-tailed test. (**D**) Schematic illustration depicting the workflow implemented to obtain and culture monocyte-derived dendritic cells (mo-DCs) with pulmonary arterial smooth muscle cells (PASMCs) from donor controls and PAH patients. Undifferentiated CD14+ cells were isolated from peripheral blood mononuclear cells (PBMCs). CD14+ cells were incubated with differentiation media to obtain immature and mature mo-DCs. Each mo-DC maturation stage was co-cultured with PASMCs in transwell at a ratio of 1:1. A separate well with only PASMCs served as a control. The image was created using BioRender. (**E**) Pulmonary arterial SMC (PASMC) proliferation (median and IQR) across co-culture conditions (immature mo-DC:PASMC; mature mo-DC:PASMC; PASMC:PASMC) and study groups (donor control and PAH patients). (**F**) Differences among culture conditions, involving immature monocyte-derived DCs (mo-DCs) and pulmonary arterial smooth muscle cells (PASMCs), for Platelet Endothelial Cell Adhesion Molecule (PECAM) and Cadherin-5 (CDH5) gene expression measured in pulmonary arterial endothelial cells (PAECs). Five technical replicates are displayed per condition. Con, donor control. Before evaluating co-culture differences, data was normalized to ‘PAEC alone’ control. Co-culture significance was assessed via a mixed-effects model, accounting for repeated measures as a random effect, followed by Tukey-adjusted pairwise comparisons. Significance levels are denoted as follows: *p < 0.05, **p < 0.01, ***p < 0.001, ****p < 0.0001.

To understand their impact on SMC proliferation, we conducted cell culture studies using mo-DCs derived from PAH and control PBMC, in combination with PASMCs from donor controls. Our results show that both immature and mature mo-DCs from PAH patients and controls induce PASMC proliferation (**Figure 4D-E**). Notably, key SMC-specific markers, including ACTA2, CXCR3, Fibronectin, SM22α, and Vimentin, remained steady or undetected (MMP9 and iNOS; **Table E3** in the Online Data Supplement). These findings align with our MIBI-TOF observations (**Figure E4D** in the Online Data Supplement). Intriguingly, the introduction of mo-DCs into the co-culture consistently reduced MMP12 gene expression in PASMCs, regardless of their origin (control or PAH-derived) (**Table E3** in the Online Data Supplement).

To delve deeper into mo-DC’s role in PASMC proliferation, we evaluated matrix metalloproteinase 12 (MMP12), and several cytokines known to impact PASMCs and derived from mo-DCs. Specifically, we examined TNF-α, IL-6, IL-1ß, IFN-α, IFN-β, IFN-γ, IL-12, and IP-10. Intriguingly, these signaling molecules were either undetected or unchanged when mo-DCs were present (**Table E4** in the Online Data Supplement), possibly because of TNF-α-induced DC maturation (21) or the 72-hour co-culture duration. Notably, mo-DCs were the source of MMP-12, while IL-6 was predominantly secreted by PASMCs (**Figure E4E**). We then assessed the influence of CD11c-DC3s on ECs, finding a consistent downregulation of PECAM and CDH5 genes in ECs, regardless of mo-DC origin (**Figure 4F**), while other genes remained unchanged (**Table E5** in the Online Data Supplement). These findings suggest that mo-DCs may contribute to the complex pathogenesis of PAH by affecting PASMC growth and influencing EC structural integrity.

### Cellular Dynamics in Hereditable PAH

In the HPAH vs. IPAH subtype comparison, we observed increased SMCs and α-SMA+ECs in the subendothelial layer of the PA (**Figure 5A**), indicating greater SMC hyperplasia and endothelial-to-mesenchymal transition (EndoMT) in HPAH patients. This corresponds with a trend toward a higher vessel score (**Figure E5A** in the Online Data Supplement), underscoring different vascular remodeling in HPAH compared to IPAH.

**Figure 5.**
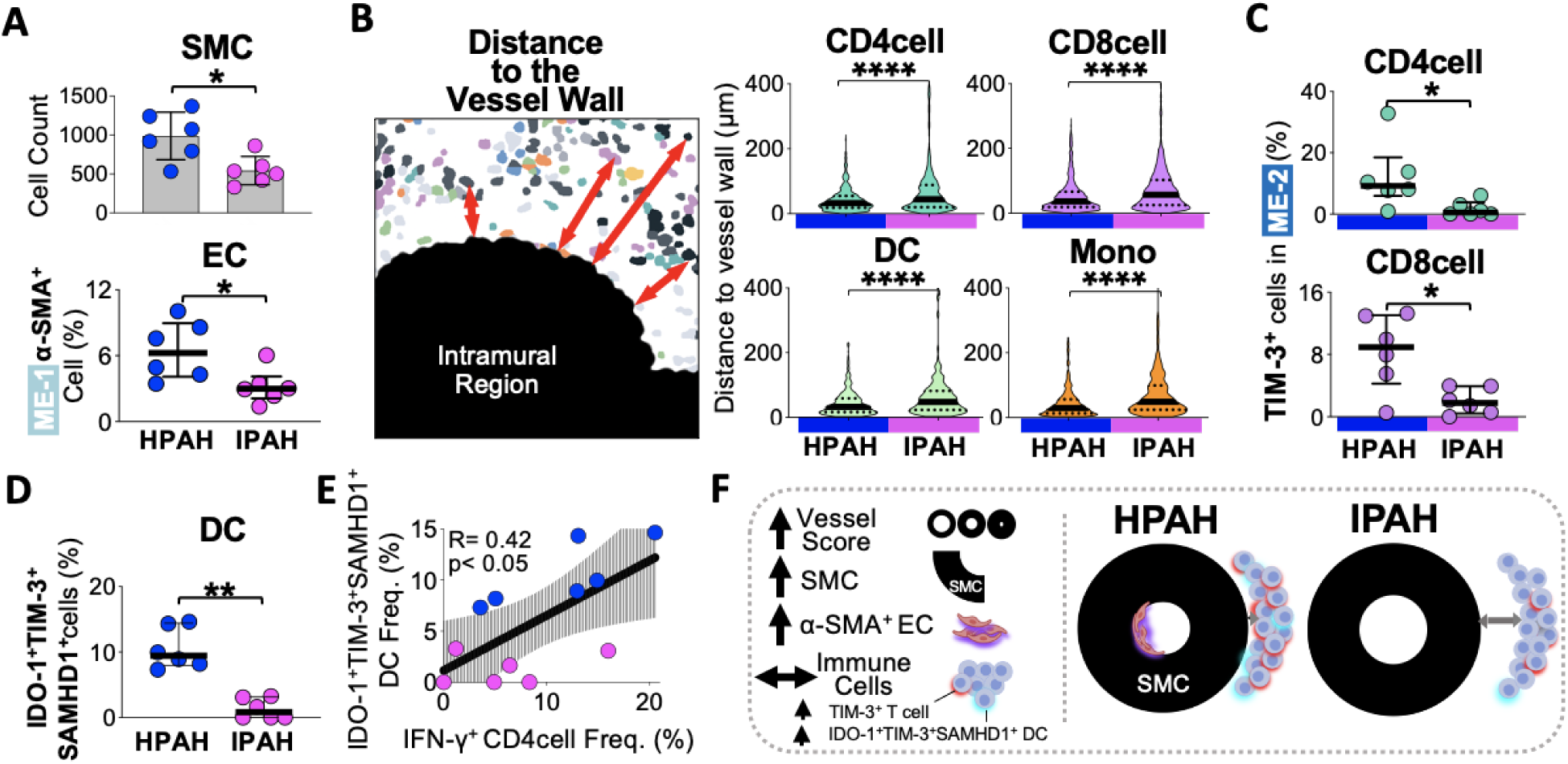
Immune dysregulation in hereditary pulmonary arterial hypertension patients. (**A**) Difference (median and IQR) in SMC counts (top) and α-SMA+ endothelial cells (EC) within ME-1 (bottom) between hereditary (H)PAH and idiopathic (I)PAH patients. **(B)** Schematic image depicting the minimum distance of extramural cell types to the intramural region (left panel). Difference in minimum distance (median, line; IQR, dotted lines) to the vessel wall of extramural immune cell subsets between PAH subtypes (right panel). **(C)** Difference (median and IQR) in TIM-3+ (CD4 and CD8) T cells within microenvironments surrounding the PA wall (ME-2) between PAH subtypes. **(D)** Difference (median and IQR) in the frequency of dendritic cells co-expressing IDO-1, TIM-3 and SAMHD1 between PAH subtypes. **(E)** Relationship (linear regression) between IFN-γ+CD4 T cells and IDO-1+TIM-3+SAMHD1+ DCs in PAH patients. Blue dots depict hereditary (H)PAH patients, while magenta dots depict idiopathic (I)PAH. **(F)** Schematic illustration summarizing the results. SMC, smooth muscle cell and fibroblast; EC, endothelial cell. P-values were calculated with a Wilcoxon rank-sum test, where *p < 0.05, **p < 0.01, ***p < 0.001, ****p < 0.0001.

Furthermore, in the HPAH subtype, extramural immune cells were closer to the vessel wall (**Figure 5B**). Among these cells, T cells dominated the periphery of PAs (ME-2), with a higher frequency of TIM-3+ T cells compared to the IPAH subtype (**Figure 5C** and **E5B-C** in the Online Data Supplement). In this dynamic microenvironment, an expanded DC subset expressing IDO-1, TIM-3, and SAMHD1 markers was noted (**Figure 5D** and **E5C** in the Online Data Supplement), exhibiting a correlation with IFNγ-producing CD4 T cells (**Figure 5E**). These findings summarized in **Figure 5F**, suggest a heightened state of immune activation and inflammation surrounding the PA wall in HPAH, potentially linking immune dysregulation to the observed cellular changes.

### Neutrophils-Endothelial Interaction in HPAH

Among various cell types, neutrophils within extramural endothelial-enriched microenvironments (ME-8) exhibited elevated levels in the HPAH subtype (**Figure 6A**). Intriguingly, neutrophil accumulation extended to immune-enriched (ME-2) and intramural endothelial-enriched microenvironments (ME-5, intima layer) (**Figure 6B**). Importantly, within the ME-5, their presence demonstrated a direct correlation with iNOS expression on PAECs and an inverse correlation with HLA-class I expression on PAECs, regardless of PAH subtypes (**Figure 6C**). However, no evidence was observed that exosomes derived from neutrophils or from other sources induce changes in PAECs (**Figure E6A-B** in the Online Data Supplement). This suggests that neutrophils may undertake distinctive functions contingent on their microenvironment, with implications for both local and systemic inflammation and potential vascular damage. Exosomes may not be a key mediator of the observed effects on PAECs from HPAH patients.

**Figure 6.**
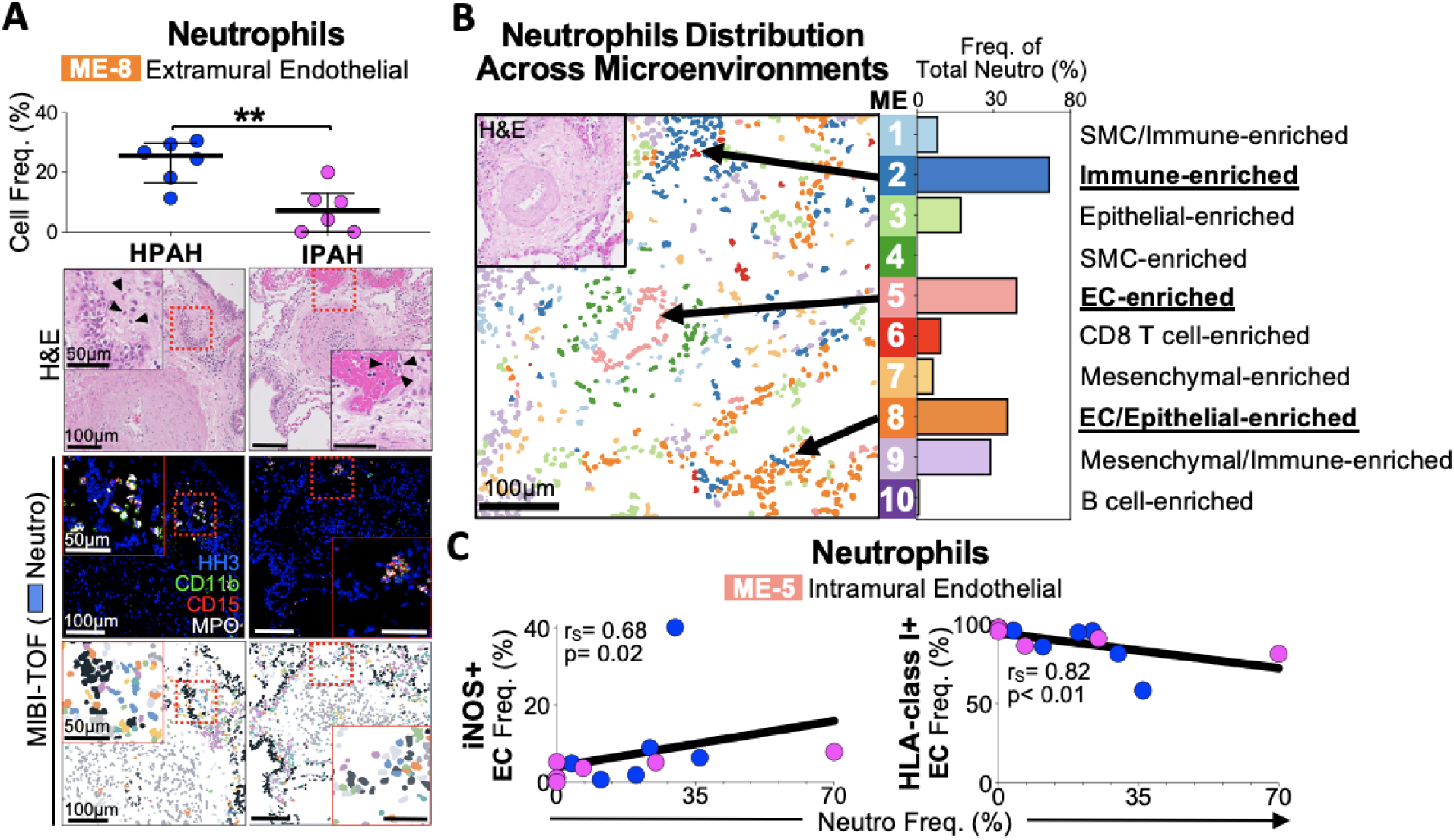
Neutrophil enrichment in hereditary PAH pulmonary artery endothelial regions. (**A**) Difference (median and IQR) in the frequency of neutrophils within extramural endothelial-enriched microenvironments (ME-8) between hereditary (H)PAH and idiopathic (I)PAH patients (top), along with corresponding representative images (bottom). Neutrophils are highlighted in the hematoxylin and eosin (H&E) zoomed insets (black arrows) and confirmed by signature markers (CD11b, CD15, MPO) using MIBI-TOF. Cell overlays from the MIBI-TOF images depict neutrophils in blue. Images scale bars = 100μm; zoomed insets scale bars = 50μm. MPO, myeloperoxidase; HH3, Histon H3. P-values were calculated with unpaired t test with Welch’s correction, where *p < 0.05, **p < 0.01, ***p < 0.001, ****p < 0.0001. **(B)** Representative image of a pulmonary artery showing the spatial localization of distinct microenvironments (MEs) on the left, along with the frequency of neutrophils across MEs on the right in PAH. Black arrows in the image indicate where neutrophils preferentially accumulate. The text highlighted in bold and underlined specifies the cell composition of these MEs. The corresponding hematoxylin and eosin (H&E) image, provides context for the PA tree architecture. Images scale bars = 100μm. **(C)** The relationship (correlation) between neutrophils within intramural endothelial-enriched microenvironments (ME-5) and the frequency of iNOS+ or HLA-class I+ endothelial cells (ECs) in patients with pulmonary arterial hypertension (PAH). Correlations were carried out using Spearman’s rank correlation (rS) with 95% confidence interval and two-tailed test, where p-values are reported as follows: *p < 0.05, **p < 0.01, ***p < 0.001, ****p < 0.0001.

## DISCUSSION

The concept of immunoprivilege within the pulmonary artery (PA) wall has been extensively investigated (22). Previous studies have primarily focused on perivascular inflammatory cells (23,24,25,26) and cytokine profiles (26,28) concerning pulmonary arteriopathies. In contrast, our previous work (5) emphasized neointimal macrophages, while this study introduces a novel finding—a correlation between intramural inflammatory cells, particularly dendritic cells (DCs), and the severity of PA occlusion. This suggests compromised immunoprivilege contributing to vascular pathology. Our results align with previous research documenting increased immune cell types in PAH (23,24,29,30,31), except for a lack of expansion in natural killer (NK) cells (32). Limited regulatory T cells (Tregs) were observed within PAH PAs, indicating an inadequate local immune control, consistent with athymic rat studies (6). Furthermore, it is intriguing to speculate that immune cells recruited in PAH may directly influence vascular remodeling through the persistent production of pro-inflammatory factors (2). Our study also revealed upregulated major histocompatibility complex (MHC) class I and II on pulmonary endothelial cells (PAECs) and smooth muscle cells (PASMCs), suggesting a mechanism amplifying intramural inflammation.

Among the various immune cell subsets, we focused on DCs, given their intramural implication in vascular pathology, and their role in modulating inflammation and shaping vascular remodeling dynamics (9,33). Our study identified a subset of DCs expressing CD14, CD209, MHC class II, and CD163, resembling monocyte-derived DCs (mo-DCs). CD163 expression, often associated with the M2 macrophage phenotype, extends to both monocytes and mo-DCs (34), including CD163+ monocytes exhibiting an inflammatory M1-like signature (35). The differentiation of mo-DCs from monocytes, under the influence of IFN-γ (36), aligns with our observation of elevated IFN-γ+ immune cells in PAH, previously documented in the PA adventitia of human and experimental PH that correlated with lesion severity (25). MIBI-TOF imaging highlighted the significant contribution of mo-DCs to vascular remodeling, independent of hemodynamic factors (25), as they accumulated within the subendothelial and adventitial regions of the PAs, mirroring the extent of lesion severity. Co-culture experiments with mo-DCs, particularly from PAH patients, revealed their ability to promote PASMCs proliferation without inducing de-differentiation.

While mo-DCs secrete various bioactive molecules, including cytokines, growth factors, and enzymes like matrix metalloproteinase-12 (MMP-12) (33, 37), the specific driver of PASMC proliferation with minimal differentiation (38) remains elusive. However, given the pivotal role of mo-DCs’ MMP12 secretion in matrix remodeling and vascular tone regulation (39), interactions with PASMC receptors may hinder MMP12 mRNA transcription in PASMCs. This could disrupt vascular homeostasis and matrix balance, potentially intensifying PAH’s complexity. Therefore, mo-DC-mediated MMP-12 impact potentially shapes vascular dynamics, extracellular matrix, and disease progression in PAH. Future studies investigating the mo-DC secretome or mo-DC-induced PASMC secretome may offer a comprehensive framework of mechanisms used by mo-DCs to influence both SMCs and ECs.

In addition to their pro-proliferative impact on PASMC, mo-DC also influence EC gene expression, by downregulating crucial EC markers involved in barrier function, such as PECAM and CDH5 (40). These alterations may affect the trans-endothelial migration of inflammatory cells in PAH (41), contributing to chronic inflammation maintenance.

Despite prior research has highlighted the heightened severity of vascular pathology in HPAH linked to a *BMPR2* mutation (26), our study unveils a novel dimension. In HPAH, a notable distinction from IPAH arises—an increase in immune cells positioned proximal to vessel walls and a prevalent presence of activated T cells. Within this dynamic microenvironment, an expanded subset of DCs akin to immune-suppressive DCs observed in rodent autoimmune models (42), emerges. These DCs correlated with IFNγ-producing CD4 T cells, suggesting a potential response to escalating inflammation, where tolerant DCs proliferate to restore immune equilibrium (43). Concurrently, the interplay between persistent inflammation and immune perturbation in PAH (44), along with reduced *BMPR2* expression, can catalyze endothelial-to-mesenchymal transition (EndoMT) (45), driving vascular remodeling and disease evolution. Our findings seamlessly align with this narrative, as the HPAH subtype exhibited heightened SMC hyperplasia and EndoMT, diverging significantly from IPAH.

Mutations in *BMPR2* and potential HPAH-related genetic variants are associated with severe disease, increased mortality (16), PA muscularization (13), and inflammation (14,15). These mutations drive the recruitment of immune cells—monocytes (46), macrophages (5), and neutrophils (15)—into the lungs, exacerbating PAH. Accumulation of neutrophils within the intima layer, correlating with increased iNOS and decreased HLA-class I expression on ECs, suggests EC damage (47) and dysfunction (48). This aligns with our prior findings of heightened neutrophil adhesion to PAECs, increased elastase production, and a tendency to form extracellular nets (41), along with myeloperoxidase, a catalyst for ROS formation (49). Importantly, our study found no evidence that exosomes derived from neutrophils or other sources induce changes in PAECs, helping narrow potential mechanisms. Additionally, HPAH patients exhibited elevated neutrophils in vessels surrounding the PAs, potentially recruiting and activating immune cells, further contributing to disease.

These findings highlight distinct roles of neutrophils influenced by their microenvironment, carrying implications for local and systemic inflammation, and potential vascular damage. Understanding this variation provides insights into disease progression, potentially guiding therapeutic strategies to mitigate neutrophil-driven vascular damage in both HPAH and IPAH.

The study’s implications for drug development in both HPAH and IPAH are significant. The presence of intramural immune/inflammatory cells, and the mo-DCs’ role in vascular remodeling and endothelial dysfunction in both PAH subtypes, emphasize the need for therapies targeting these features. In HPAH, immune dysregulation, increased neutrophils, and α-SMA+ECs highlight the demand for additional targeted strategies.

While our study offers valuable insights, certain limitations warrant consideration. The use of lung tissues from end-stage PAH patients might confound inflammation with tissue injury, oxidative stress, and metabolic dysfunction. Systemic prostacyclin therapy in PAH patients, known to influence inflammation (50), may introduce inflammation-associated signatures and drug-related effects, potentially masking primary pathogenic signals, and early events. Prospective research could harness advanced profiling methods, like integrated single-cell spatial transcriptomics and protein mapping for a deeper understanding of PAH’s immune dysregulation, aiding in innovative immunotherapies. While broadening the study to various PAH stages could mitigate end-stage variations, logistical challenges in obtaining tissue samples make this approach unfeasible. Consequently, our focus remained on studying vessels at different pathological occlusion stages through lung transplant tissue.

In conclusion, our study uncovers immune infiltration in vessel walls, disrupting pulmonary arterial immunoprivilege and correlating with the severity of PA occlusive changes. We identify TIM-3+ T cells and IDO-1+TIM-3+SAMHD1+ dendritic cells as novel contributors to immune dysregulation driving PAH progression. Notably, mo-DCs significantly impact vascular remodeling and endothelial dysfunction, offering promising therapeutic avenues. These findings advance PAH understanding, paving the way for improved treatments and patient outcomes.

### Data and Code Repository

All custom code used to analyze data will be made available through a Github repositories (https://github.com/angelolab and https://github.com/SelenaFerrian) and all processed images and annotated single cell data will be made available in Mendeley’s data repository.

