## Supplementary material for "Single-Cell Imaging Maps Inflammatory Cell Subsets to Pulmonary Arterial Hypertension Vasculopathy": Online Data Supplement

### EXTENDED METHODS

#### Study Cohort

Lung tissues at the time of transplant from PAH and unused donor controls (Con) were provided by the Pulmonary Hypertension Breakthrough Initiative (PHBI), funded by the NIH-NHLBI grant R24 HL123767, and the Cardiovascular Medical Research and Education Fund (CMREF) UL1RR024986. The tissues were procured under IRB approval at the Transplant Procurement Centers of the PHBI, and de-identified patient data were obtained via the Data Coordinating Center at the University of Michigan. The study cohort represented 22 patients: Con (n=10), HPAH (n=6) and IPAH (n=6). Among the inclusion criteria for the unused donor controls, we selected lungs with no signs of infiltrates on the chest x-ray taken within 24 hours of donor's death, absence of fever >38.2°C (100.76°F) within 24 hours of donor's demise and lungs with unexpected pulmonary problems such as significant emphysema or pulmonary fibrosis noted on radiographic studies or significant pulmonary hypertension. Controls were age- and gender-matched to the PAH patients. All HPAH patients, except for PAH05 who had no delineated mutations but a significant family history of PAH, carried either a mutation in the *BMPR2* gene or the *SMAD9* gene. Clinical data are summarized in **Table 1**. All PAH patients were on triple vasodilator therapy. This included sildenafil, a prostacyclin analog (either epoprostenol, iloprost, treprostinil or combinations of those) and an endothelin receptor antagonist (either bosentan, ambrisentan, and/or sitaxsentan). One patient also received the tyrosine kinase receptor inhibitor (imatinib). Peripheral blood mononuclear cells (PBMCs) were isolated from PAH and donor control blood obtained from the Stanford biobank and the Stanford Blood Bank, respectively.

### Histopathological Classification of Pulmonary Arteriopathies

PAH and control samples showing impaired inflation or extreme hemorrhage that obscured vascular pathology were excluded. PAH samples with evidence of veno-occlusive disease were also excluded. Similarly, Con samples with no signs of inflammation or significant vascular change were included. Serial sections of each specimen were stained with hematoxylin and eosin (H&E) for region selection. Approximately five representative regions of interest (ROIs) were selected per patient based on scoring of vascular morphology as performed by two co-authors (TS, MR) (**Figure E1A**). Two $mm^2$ core tissue was embedded in a tissue microarray (TMA). To standardize pathological reporting and to provide a simplified description of the severity of each vascular lesion, we recorded the extent of medial and neointimal thickening or whether the lesion was plexogenic and the size of the artery affected focusing on those at terminal and respiratory bronchiolus levels (**Figure E1B**). Scores were designated as follows: Normal (Score 0) – medial wall thickness ≤20% of total vessel diameter which included some vessels with a mild neointima; medial hypertrophy (Score 1) – medial wall thickness including mild neointimal formation resulting in between >20 and <50% of lumenal occlusion; mild neointimal lesions (Score 2) – lumenal occlusion in between ≥50% and <75% of total vessel diameter; moderate neointimal lesions (Score 3) – lumenal occlusion ≥75% but <90% of total vessel diameter; severe neointimal lesions (Score 4) – lumenal occlusion ≥90% to complete occlusion; plexiform lesion (Score 5) – complete lumenal occlusion associated with an aberrant angiogenic response.

### Gold Slide Preparation

Slide preparation was performed at the Stanford Nano Shared Facility (SNSF), as previously described (17). Prior use, gold-coated slides were silanized in acetone with 3-

aminopropyltriethoxysilane for 30 min then washed with acetone and air-dried (Vactabond, Burlingame, CA). Slide were baked at 70°C for 30 min and kept dry at room temperature.

**MIBI-TOF Reagents Quality Control Prior Labeling**

Prior to metal conjugation, each antibody was screened with a NanoDrop spectrophotometer using a mouse IgG standard curve. Protein concentration and purity were recorded to assure optimal metal conjugation. Following nanodrop measurements, SDS-PAGE electrophoresis was carried out according to the manufacturer instructions (Bio-Rad Laboratories) to test antibodies for carriers and protein degradation that could interfere with metal-labeling. Precast SDS gels were imaged using Alpha Innotech AlphaImager 2200 Imaging System (Thermo Fisher Scientific). Acceptable quality IgG or IgM sample showed typical high and low molecular weight bands as seen in control lane and absence of bovine serum albumin (BSA) or other carriers.

**Tissue Microarrays Preparation**

Three tissue microarrays (TMAs) were generated. Appropriate positive and negative tissue controls were selected and compiled into a 'titration TMA' to assess the specificity and sensitivity for each target with regard to its cellular localization, subcellular localization, and co-expression of proteins across different tissue types. These included spleen, tonsil, lymph node, placenta, colon, normal lung and diseased lung tissue from three PAH patients. The titration TMA was used to screen each antigen by single-plex chromogenic IHC and MIBI-TOF staining. Additional control tissues comprising normal lymph node, spleen, and tonsil were compiled into the study 'cohort TMAs' to assure intra- and inter-assay precision across MIBI-TOF acquisitions.

**Antibody-Labeling**

A summary of antibodies, clone, reporter isotopes and provider information can be found in **Table E1**. Antibodies were conjugated to isotopic metal reporters as described previously (17). Following conjugation, metal-antibody affinity was assessed by ICP-mass spectrometry to assure a total recovery of 80-100 atoms per antibody. Antibodies were then diluted in Candor PBS Antibody Stabilization solution (#130050, Candor Bioscience) to 0.2 mg $mL^{-1}$. Antibodies were lyophilized in 100 mM D-(+)-Trehalose dihydrate (#T9449, Sigma Aldrich) with ultrapure distilled $H_2O$ for storage at 20°C. Prior to staining, lyophilized antibodies were reconstituted in a buffer of Tris (#15567027, Thermo Fisher Scientific), sodium azide (#S2002, Sigma Aldrich), ultrapure water (#10977015, Thermo Fisher Scientific), and antibody stabilizer to a concentration of 0.05 mg/mL.

**Antibody Titration and Validation**

Three serial sections of the 'titration TMA' were placed on the same gold slide to identify the working titer for MIBI-TOF. Six serial dilutions were performed for each antibody to identify the working titer that yielded maximal separation between signal and noise across tissues. When a negative tissue control was unavailable, appropriate fields of view were evaluated, as previously described (E1). Accuracy of MIBI phenotyping was assessed by comparison to conventional single-plex chromogenic IHC on the same TMA to ensure target specificity and sensitivity across tissue types (**Figure E1F**).

**Antibody Panel and Staining**

A total of 90 ROIs were selected as being in the same section as a PA lesion and scanned using MIBI-TOF. Each ROI was approximately 500 $mm^2$ and 1024 x 1024 pixels with a resolution of ~ 500 nm. Single sections (4 μm-thick) were obtained for histological and multiplexed immunohistochemical examination. To ensure target specificity and sensitivity, clinical FFPE tissue specimens in parallel with tissue controls for each marker were placed on gold slides and stained overnight using a cocktail single master mix comprising 35 metal-labeled primary antibodies (**Figure E1C**). All targets were stained together without the need for cyclical steps or enzymatic reactions (17). The full panel across its respective tissue control is reported in **Figure E1F**. Such a panel included targets to identify lineage cell types, immune infiltrate, and functional proteins associated with inflammatory immune and vascular cell responses (**Table E1**). Staining was performed as previously described (17). Briefly, slides were baked at 70$^{o}$C for 20 minutes in the oven. De-paraffinization and rehydration was carried out in Xylene (X3), 100% EtOH (X2), 95% EtOH (X2), 80% EtOH, 70% EtOH and dd$H_2O$ (X2) using the Leica ST4020 Small Linear Stainer (processing time= 30 sec; number of dips= 3). Sample were transferred to the 75$^{o}$C preheated 1:10 Dako Target Retrieval Solution pH 9 (3in1) (#S2375, Dako) using a Potential Transformer (PT) module (#A80400012, Thermo Scientific) filled with 1.5L of 1X low-Barium PBS. Heat induced epitope retrieval was carried out at 97$^{o}$C for 40 minutes. Once the heating cycle was finalized, sample were cooled to 65$^{o}$C for 20 minutes within the PT module, followed by room temperature for 15 minutes. Prior protein blocking, a washing step (X2) in 1:20 wash buffer (#935B-09, Cell Marque) supplemented with 0.1% BSA was carried out for 5 minutes. BBDG Blocking Buffer (2% Normal Donkey Serum, 1% BSA, 0.1% Cold Fish Skin Gelatin, 0.1% Triton X-100, 0.05% Sodium Azide, resuspended in 1X TBS IHC Wash Buffer with Tween 20 in dd$H_2O$) was implemented to prevent non-specific binding. Samples were incubated with BBDG Blocking buffer for 1 hour at room temperature. An endogenous Biotin Blocking (#927301, Biolegend) step was introduced for biotinylated antibodies included in the antibody cocktail (PD-L1b). Samples were treated with Avidin for 10 minutes at room temperature, followed by a washing step (X2) for

5 minutes and a final incubation with Biotin for 10 minutes at room temperature, followed by another washing step for 5 minutes. The *antibody cocktail 1* was prepared in 1:20 of TBS IHC Wash Buffer with Tween 20 (#934B-09, Cell Marque) supplemented with 3% NDS and filtered through a 0.1 μm centrifugal filter (#UFC30VV00, Millipore) at 12,000 x g for 2 minutes. The antibody mix was added to the samples, after removing BBDG blocking buffer, for overnight incubation at 4°C in a sealed humidity chamber. The following day, samples were washed twice for 5 minutes in wash buffer and the *antibody cocktail 2* was prepared as previously and incubated with the samples for 1 hour at 4°C. Following incubation, slides were washed twice for 5 minutes in wash buffer and incubated for 5 minutes with a fixative solution of 2% glutaraldehyde (#50-262-13, Electron Microscopy Sciences) in low-barium PBS. Slides were washed three times in PBS (X1), 0.1 M Tris at pH 8.5 (X3), dd$H_2O$ (X2), and then dehydrated by washing in 70% EtOH, 80% EtOH, 95% EtOH (X2), 100% EtOH (X2). Prior to MIBI-TOF imaging, slides were dried under vacuum for at least of 3 hours.

**MIBI-TOF Imaging**

Imaging was performed using a MIBI-TOF instrument with a Hyperion ion source. $Xe^+$ primary ions were used to sequentially sputter pixels for a given field of view (FOV). A total of 3330 images from the cohort and 370 from the tissue controls were generated using the following imaging parameters: field size, 500 $\mu m^2$ at 1024 × 1024 pixels; dwell time, 4ms; ion dose per unit area, 442 nAmp hours / $mm^2$ for 500 $\mu m^2$ FOVs; median gun current on tissue: 1.36 nA $Xe^+$.

**Image Processing And Single-Cell Segmentation**

Image processing and nuclear segmentation were carried out as previously described (17) (**Figure E1D-E**). Briefly, multiplexed images were extracted, background subtracted, denoised, and aggregate filtered. Following image processing, nuclear segmentation was performed using an adapted version of the FeatureNet convolutional neural network (E2). The nuclear channel was used to identify nuclei in each image. To train a neural network capable of accurately segmenting these images, 8 distinct fields of view were manually labeled using the DeepCell Label annotation software (https://github.com/vanvalenlab/deepcell-label). Pan-CK and Na+K+-ATPase were used to visualize the cell membrane to guide manual segmentation of training data. Following manual labeling of each cell, segmentation masks were transformed into binary masks marking the nuclear interior, nuclear border, and non-nuclear background. The images were cropped into tiles of 128x128 pixels, and underwent extensive augmentation such as rotations, crops, flips, and scaling. The augmented data was split into a train and validation split for network training. Following prediction of the nuclear borders, segmentation predictions were expanded by a 3-pixel radius to approximate the boundaries of each cell.

**Single-Cell Phenotyping and Composition**

Cell clustering analyses were carried out using pre-processed (background subtracted and de-noised) images as previously described (18). Single cell data were extracted for all cell objects and normalized to cell area. Cells with a sum of less than 0.1 normalized to total area counts across all lineage channels were excluded from analysis. Single cell data was linearly scaled with a scaling factor of 100 and arcsinh-transformed with a co-factor of 5. All mass channels were scaled to 99.9th percentile. In order to assign each cell to a lineage, the FlowSOM clustering algorithm was used in iterative rounds with the Bioconductor "FlowSOM" package in R (19). The first clustering round separated cells into 100 clusters that were subsequently merged into five major cell lineages using the "Metaclustering_consensus" function: immune (CD45+Vim+), epithelial (Pan-CK+Vim+), mesenchymal (Vim+), endothelial (CD31+Vim+), and SMC (α-SMA+Vim+ and α-SMA+Vim-) which includes both SMCs and fibroblasts. Proper lineage assignments were ensured by overlaying FlowSOM cluster identity with lineage-specific markers. No cell remained unclassified using the combination of these five markers. Immune cells were then subclustered again to delineate B cells (CD20+), CD4+ T cells (CD3+CD4+), CD8+ T cells (CD3+CD8+), Tregs (CD3+CD4+FOXP3+), NKs (CD14-CD16+), neutrophils (CD15+CD11b+) and mononuclear phagocytes (macrophages, monocytes, and dendritic cells) using a combination of myeloid markers such as CD68, CD14, CD11c, CD11b, CD16, DC-SIGN (**Figure 1B**). The monocyte population, characterized as CD14+CD209-CD163-CD68-, encompassed both monocytes and fibrocytes (α-SMA+). The relative abundance of all major lineages was determined out of total cells per FOV and the relative frequency of immune cell subsets was determined out of total immune cells per FOV. DCs, were further subclustered to delineate different DC subsets using CD14, CD11c, CD141, CD209, CD11b, CD163, and HLA-DR markers. Four DC subsets were identified (**Figure E4A**) as CD11c+ DC1s, CD11c+ DC2s, CD11c- DC3, and CD11c- DC4.

**Vascular Region Masking**

In order to assess the localization of immune cells in PAH patients with respect to vascular architecture, binary masks of the intramural region (IM) were computationally generated for each FOV using the α-SMA staining (**Figure 2A**). A region within 100 px (~50 μm) of the vessel was defined as a 'perivascular' (PV) zone. Any region beyond that was considered 'non-vessel' (NV). Masks were further processed in Matlab to fill any holes, exclude objects smaller than 1000 pixels, and dilate the mask to smooth edges. Next, cells were assigned to belong to the IM region if they had complete overlap with the binary mask. Cells on the mask boundary or outside of the mask were designated as PV, while cells outside any of these masks were designated as NV.

**Functional Analyses**

Positivity thresholds for functional proteins were customized to each marker individually using a silhouette scanning approach from the MetaCyto software in R (20). The point separating the two peaks, representing the positive and negative (background) marker expression on a bimodal distribution, was used as cut-off value for determining positivity of a marker. Thresholds were further validated by visual inspection of positive and negative cells in image sets (18).

The co-expression of functional markers on DCs was further examined using a Boolean combinations approach, to evaluate all possible combinations of individual markers partitioned into positive/negative categories. This allowed us to identify DCs that expressed antigen ‘x’ and ‘y’, but not ‘z’, based on the ‘AND’, ‘OR’ and ‘NOT’ logic.

**Spatial Analyses**

The intercellular relationship results in an alteration of the tissue architecture, which we aimed to capture through a spatial analysis. To analyze the cellular associations with the pulmonary arteries, the minimum distance for each immune cell centroid to the nearest perimeter location of the intramural mask (described above) were calculated, thus identifying which immune cell types were prone to locate in proximity to the vessel.

To visualize and quantify cell-cell-neighborhood relations, the information on cell type and position was used to phenotype local neighborhoods and reveal how their spatial distribution leads to the generation of tissue architecture (E3). To accomplish this, cell neighborhoods were produced by first generating a cell neighbor matrix, where each row represents an index cell, and the columns indicate the relative frequency of each cell phenotype within a 25 μm (50 pixel) radius of the index cell. On average, this algorithm picked up the cell itself and the initial 10-13 cells that this cell was interacting with, thus grouping cells into local microenvironments (**Figure 3A** and **E3A**). Next the neighbor matrix was clustered into 10 clusters using k-means clustering. We determined the optimal number of clusters (k) through a combination of quantitative and expert manual curation.

Initially, the elbow method suggested k=5, but upon visual inspection, we observed that it did not capture the necessary details, such as tertiary lymphoid structures (TLSs). We then experimented with different k values, including k=6, 7, 8, 9, and 10, aiming for a balance between granularity and biological relevance. Ultimately, k=10 emerged as the optimal choice, effectively identifying TLSs, and aligning well with known cell types and tissue structures. The generated neighborhoods contain information on the cell composition, density, expression of specific molecules, and data on any additional structural or functional parameters (**Figure 3B-C**). Neighborhood cellular profile was determined by assessing the mean prevalence of each cell phenotype in a 25 μm radius of all index cells assigned to that neighborhood, while functional marker expression was determined by assessing mean marker expression by the index cells assigned to each neighborhood cluster.

**Pulmonary Arterial Smooth Muscle Cell and Endothelial Cell Growth**

Primary human pulmonary arterial SMCs (PASMCs) from a donor control (Con) patient were harvested from small pulmonary arteries <1 mm in diameter from unused donor lungs. Briefly, the smooth muscle layer of PAs was digested in 1 mg/mL collagenase (#C9722, Sigma) and 75 μg/mL elastase (#2292, Worthington) in HBSS (#14025092, Thermo Fisher Scientific) at 37$^{o}$C for 25 min as described before with modification (E4). PASMC were spun down at 1,200 RPM for 5 min and seeded in T25 flasks with SMC growth medium (Smooth Muscle Growth Medium-2 containing 5% fetal bovine FBS #CC-3182, Lonza). PASMCs were subcultured at a 1:3 ratio and used between passages 3-5.

PAECs (#C-12241, PromoCell) were cultured in EC medium supplemented with 5% FBS, 1% P/S, and 1% ECGS (#1001, ScienCell Research Laboratories), and used between passages 4 and 7.

**Generation of Monocyte Derived-Dendritic Cells**

The workflow for the mo-DCs generation and co-culture is depicted in **Figure 4D**. Peripheral blood mononuclear cells (PBMCs) were isolated using Ficoll-Hypaque (#H8889, Sigma Aldrich) by density gradient centrifugation. Human CD14+ cells from a donor control (Con) and an IPAH patient's PBMC were obtained using human CD14 Microbeads (#130-096-53, Miltenyi). CD14+ monocytes were differentiated into immature human mo-DCs by culturing for 7 days using CellXVivo Human Monocyte-derived DC Differentiation Kit (#CDK004, R&D Systems) according to the manufacturer protocol. Briefly, on day 0 freshly isolated monocytes were supplemented with Differentiation Media containing recombinant human IL-4 and GM-CSF to generate immature mo-DCs and incubated for 3 days in a humidified incubator at 37°C and 5% $CO_2$. On day 3, half of the media was removed and replenished with the same volume of fresh Differentiation Media. Cells were further incubated for an additional 2 days. On day 5, the process of changing the media was repeated and cells were incubated for an additional 2 days. On day 7, immature mo-DCs were observed and confirmed by flow cytometry. Along with IL-4 and GM-CSF, a portion of immature mo-DCs were supplemented with recombinant human TNF-α and incubated for additional 3 days to obtain mature mo-DCs. On day 10, mature mo-DCs were observed under microscope. Phenotypic analysis was conducted on day 7 for immature mo-DCs and on day 10 for TNF-α treated mature mo-DCs. Cells were stained with the indicated antibodies for DC-SIGN/CD209 (#330113, Biolegend), B7-1/CD80 (#564160, BD Biosciences), B7-2/CD86 (#562390, BD Biosciences), CD163 (#333615, Biolegend), CD14 (#347493, BD Biosciences), HLA-DR (#307606, Biolegend), or an appropriate isotype control antibody. Stained cells were analyzed using a LSRII Fortessa.

**Monocyte Derived-Dendritic Cells and Vascular Cells Co-Culture**

To address the interaction between vascular cells and mo-DCs we conducted *in vitro* co-cultures (**Figure 4D**). Immature and mature mo-DCs were cultured with PASMCs or PAECs using 0.4 µM pore size transwells (#CLS3470, Sigma Aldrich) in a 1:1 ratio. PASMCs were seeded at the bottom of the transwell in SMC growth medium for overnight followed by 24 hours starvation in 0.2% FBS SMC medium before co-culture. PASMC or mo-DCs were seeded on the insert filter of the transwell and incubated in 5% FBS SMC medium for 72 hours. The co-cultures were performed in humidified incubator at 37°C and 5% $CO_2$. Following co-cultures, SMCs were lift by TrypLE express (#12605010, Thermo Fisher Scientific) and used for next step. Four to six technical replicates for each measurement were produced. The same steps were applied to mo-DCs and PAECs co-cultures, except for the utilization of EC culture medium. After co-culturing, PASMCs were detached by TrypLE express and counted (Scepter).

The co-culture experiments were replicated two times, with different patients and donors, using immature mo-DCs and the method described above, with slight modifications: PASMCs viability and proliferation were assessed by CellTiter 96® $AQ_{ueous}$ One Solution Cell Proliferation Assay (MTS assay) (MTS) assay (#G3582, Sigma Aldrich) following the kit instruction. Four to six technical replicates for each measurement were produced.

**Exosome Isolation from Plasma and CD66b Pulldown**

Exosome isolation from plasma and CD66b pulldown were performed as previously outlined (41). Briefly, human plasma (2 mL) from three control individuals (Con) and six PAH patients was processed. Total extracellular vesicles (EVs) were initially enriched through a series of differential centrifugation steps (10 min at 300 g, 10 min at 2000 g, 30 min at 10,000 g, and 70 min at 100,000 g, all at 4°C) using a Sorvall WX+ Ultracentrifuge (Thermo Fisher Scientific).

Total exosomes were subsequently purified using CD66b/CEACAM 8 antibodies (#SAB4301144, Sigma Aldrich) as a specific marker for human neutrophils. These antibodies were conjugated to beads using the Dynabeads Antibody Coupling Kit according to the manufacturer's instructions (#14311D, Thermo Fisher Scientific). The exosomes were co-incubated with magnetic beads conjugated with CD66b/CEACAM 8 antibodies on a shaker for 18 hours at 4°C. After incubation, unbound exosome supernatant was removed using a magnet, and the captured beads were washed.

To dissociate the beads from the CD66b/CEACAM 8-positive exosomes, an acid wash was performed by adding 200 μL of a 50 mM pH 2.5 glycine solution to the tube containing the beads for 10 minutes. Subsequently, 70 μL of a 1M pH 7.5 Tris buffer was added to neutralize the solution. The concentration of the eluted CD66b-positive exosomes and the unbound CD66b-negative exosomes was quantified using the Fluorocet Ultrasensitive Exosome Quantitation Assay Kit (#FCET96A-1, System Biosciences).

**Exosomes and Endothelial Cells Co-Cultures**

To study PAEC-exosome interaction, we conducted *in vitro* co-cultures (**Figure E6A**). A total of 5000 PAECs (#C-12241, PromoCell) were initially seeded on 96 well plates and cultured overnight in humidified incubator at 37°C and 5% $CO_2$ using EC medium supplemented with 5% FBS, 1% P/S, and 1% ECGS (#1001, ScienCell Research Laboratories), followed by 24 hours serum starvation. An optimal dose of exosomes (10,000 count/cell) from both CD66b+ and CD66b- sources were applied to PAECs using 0.4 μM pore size transwells (#CLS3470, Sigma Aldrich) in a 1:1 ratio. Co-cultures were maintained for 72 hours in a humidified incubator at 37°C and 5% $CO_2$.

**Real-time PCR assessed mRNA expression**

SMC and EC total RNA was extracted using QiAzol (#79306, ZyQiagen). The RNA's quantity and quality were assessed using a spectrophotometer (Biotek Synergy H1 hybrid reader). Reverse transcription was carried out using the High Capacity RNA to cDNA Kit (#4374967, Applied Biosystems). RT-qPCR was performed by using PowerUp SYBR green PCR Master Mix (#A25778, Applied Biosystems) and conducted in triplicate using a CFX384 Real-Time System (BioRad), following the procedure described previously (E5). The relative transcriptional level of mRNA was calculated using the $2^{-\Delta\Delta Ct}$ relative quantitative method, where $\Delta\Delta Ct = \Delta Ct$ (experimental group) - $\Delta Ct$ (normal group), $\Delta Ct = Ct$ (target gene) - Ct (internal reference), and $2^{-\Delta\Delta Ct}$ represented the relative mRNA transcriptional level. Expression levels of the selected genes (B2M, PECAM1, CDH5, ACTA2, Snail, Slug, CXCR3, CD14, MMP9, MMP12, Vimentin, SM22a, Fibronectin, iNOS) were normalized to B2M. Four to six technical replicates were produced for each measurement. The primer sequences were designed using NCBI's Primer-BLAST function or PrimerBank (http://pga.mgh.harvard.edu) and are listed below:

| | Forward Primer | Reverse Primer |
|---|---|---|
| B2M | 5'-TTCTGGCCTGGAGGCTATC-3' | 5'-TCAGGAAATTTGACTTTCCATTC-3' |
| PECAM1 | 5'-GCAACACAGTCCAGATAGTCGT-3' | 5'-GACCTCAAACTGGGCATCAT-3' |
| CDH5 | 5'-GTTCACCTTCTGCGAGGATA-3' | 5'-GTAGCTGGTGGTGTCCATCT-3' |
| ACTA2 | 5'-CCCTGAAGTACCCGATAGAACA-3' | 5'-GGCAACACGAAGCTCATTG-3' |
| Snail | 5'-CGAGTGGTTCTTCTGCGCTA-3' | 5'-CTGCTGGAAGGTAAACTCTGGA-3' |
| SLUG | 5'-TGGTTGCTTCAAGGACACAT-3' | 5'-GTTGCAGTGAGGGCAAGAA-3' |
| CXCR3 | 5'-ACGAGAGTGACTCGTGCTGTAC-3' | 5'-GCAGAAAGAGGAGGCTGTAGAG-3' |
| CD14 | 5'-ACGCCAGAACCTTGTGAGC-3' | 5'-GCATGGATCTCCACCTCTACTG-3' |
| MMP9 | 5'-TGGCTTTTGTGACAGGCACTTC-3' | 5'-CGGTGGTGTTCTCCAATGTAAGAG-3' |
| MMP12 | 5'-TGGCCATTCCTTGGGGCTGC-3' | 5'-GGGGGTTTCACTGGGGCTCCATA-3' |
| Vimentin | 5'-AGTCCACTGAGTACCGGAGAC-3' | 5'-CATTTCACGCATCTGGCGTTC-3' |
| SM22-alpha | 5'-CCGTGGAGATCCCAACTGG-3' | 5'-CCATCTGAAGGCCAATGACAT-3' |
| Fibronectin | 5'-AGGAAGCCGAGGTTTTAACTG-3' | 5'-AGGACGCTCATAAGTGTCACC-3' |
| iNOS | 5'-GCCAGGCCACCTCTATGTTT-3' | 5'-ATAGCGCTTCTGGCTCTTGA-3' |
| PECAM1 | 5'-GCAACACAGTCCAGATAGTCGT-3' | 5'-GACCTCAAACTGGGCATCAT-3' |

#### Enzyme-linked immunosorbent assay (ELISA)

Cytokines were quantified in the cultured media using ELISA following the manufacturer's guidelines. We measured human TNF-alpha (#DY210, R&D Systems), human IL-6 (#DY206, R&D Systems), human IL-12 p70 (#DY1270-05, R&D Systems), human CXCL10/IP-10 (#DY266-05, R&D Systems), human IFN-gamma (#DY285B-05, R&D Systems), human IFN-beta (#DY814-05, R&D Systems), human IFN-alpha/IFNA2 (#DY9345-05, R&D Systems), and MMP12 (#EH327RB, Thermo Fisher Scientific).

#### Software And Statistical Analyses

Image processing was conducted with Matlab 2019a. Statistical analysis was conducted in Matlab 2019a, and R v3.6.0. Data was first tested for normality with the Shapiro-Wilk test and D'Agostino & Pearson test. Normally distributed data was compared between two groups with the two-tailed Welch's T-test. Nonparametric data was compared using the Mann–Whitney U Test. Spearman's rank correlation and ordinal logistic regression were utilized to evaluate the strength and direction of the relationship between predictors and vessel score. Prior to comparing differences among the co-culture conditions, the data was normalized relative to the PASMC or PAEC controls. Significance among the co-culture conditions was assessed using a mixed-effects model that accounted for within-condition repeated measures as a random effect, followed by pairwise comparisons between the estimated marginal means (EMMs), with Tukey adjustment applied to control the family-wise error rate. Results were expressed as estimate, standard error (SE), t ratio and p value, where ns $p > 0.05$, *$p < 0.05$, **$p < 0.01$, ***$p < 0.001$, ****$p < 0.0001$. Representative images were processed in Adobe Photoshop CC2019. Graphs were produced with R v3.6.0 and Prism v10.0.0, while schematic visualizations were produced with Biorender.

### SUPPLEMENTARY FIGURES

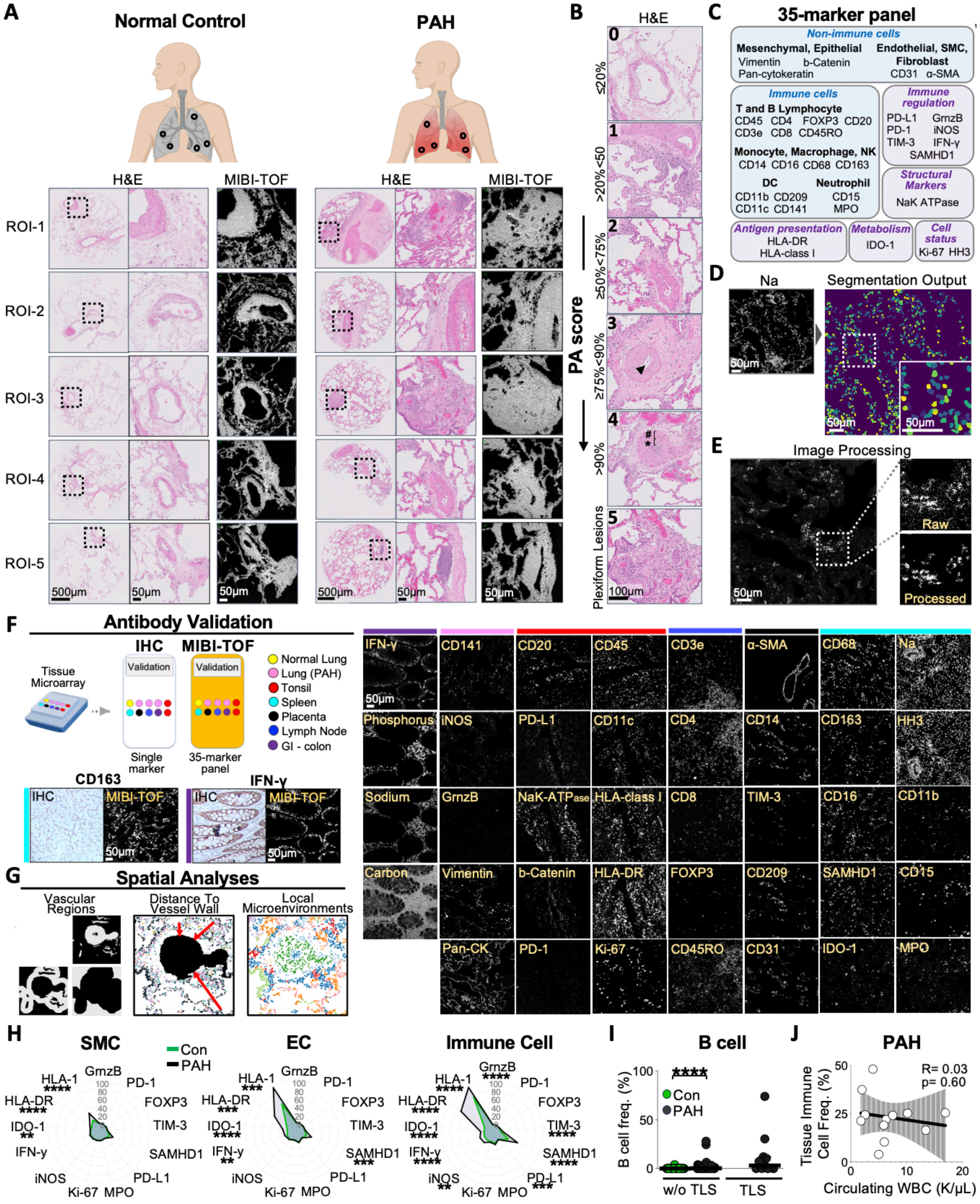

**Supplementary Figure E1. Pathology and immune infiltration in PAH: MIBI-TOF analysis.** **(A)** Overview of sample selection. Paraffin embedded lung tissues selected for the presence of at least one pulmonary artery (PA), at the level of a terminal or respiratory bronchioles. Serial sections of each specimen were stained with hematoxylin and eosin (H&E) for region selection and imaged using MIBI-TOF. Approximately five ROIs were selected per patients. H&E image scale bars = 500 µm; zoomed inset scale bars = 50 µm. **(B)** Representative hematoxylin and eosin (H&E) images of PAs from PAH patients scored from 0 to 5 according to the severity of the vascular pathology, characterized by medial hypertrophy (hash symbol), neointimal formation and fibrosis (asterix symbol) causing varying degrees of lumenal occlusion (arrow). See Methods section. Scale bars = 100 µm. **(C)** Overview of the MIBI-TOF panel. Markers were grouped by target cell type or protein class. **(D)** Workflow for Deepcell-based nuclear segmentation of single cells from multiplexed images. Na, Sodium. Image scale bars = 50 µm. **(E)** Workflow for image processing. Multiplexed images were extracted, background-subtracted, denoised, and aggregate-filtered as previously described (21). Images scale bars = 50 µm. **(F)** MIBI-TOF antibody validation was conducted across various tissue types to assess the antibody's specificity and effectiveness. This was contrasted with single-plex immunohistochemistry (IHC) to evaluate the accuracy and efficiency of the MIBI-TOF method. GI, gastrointestinal. Images scale bars = 50 µm. **(G)** Schematics of spatial analyses conducted, showing the positioning of cells in relation to vascular regions, their proximity to the vessel wall, and the intercellular distances, thus establishing distinct local neighborhoods. **(H)** Difference in functional marker expression between donor control (Con, green line) and pulmonary arterial hypertension (PAH, black line) patients in smooth muscle cells (SMC), endothelial cells (EC), and immune cells. The median frequency of positive cells per marker and study group is displayed. **(I)** Difference in B cell frequency in pulmonary PAs with or without tertiary lymphoid structures (TLS) between donor control (Con, green) and pulmonary arterial hypertension (PAH, black) patients. **(J)** Relationship (linear regression) between the circulating white blood cell count (WBC) and tissue immune cell infiltration in patients with PAH. The p-values are reported as follows: *p < 0.05, **p < 0.01, ***p < 0.001, ****p < 0.0001. GrnzB, Granzyme B; MPO, Myeloperoxidase; HLA-I, human leukocyte antigen class I; HLA-DR, human leukocyte antigen – DR isotype; HH3, Histon H3; α-SMA, alpha-smooth muscle actin; DC, dendritic cell; Mono, monocyte; Macro, macrophage; Neutro, neutrophil; NK, natural killer (NK) cell. The image was created using BioRender.

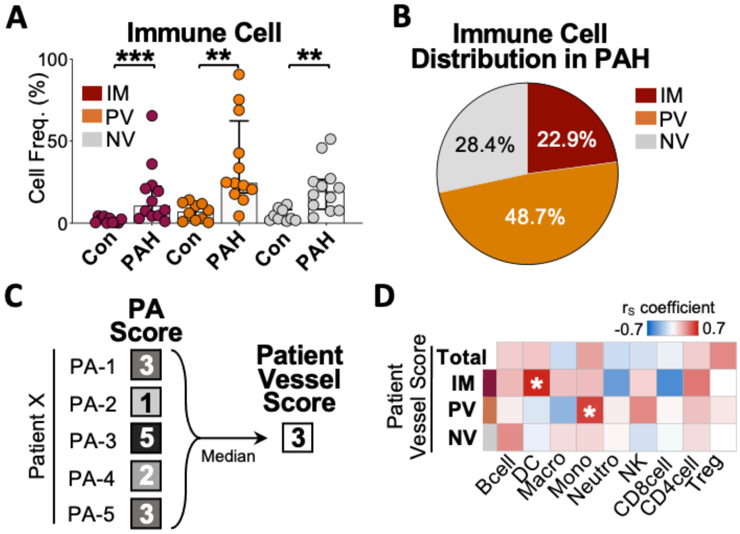


**Supplementary Figure E2. Immune cell distribution and vessel scores in PAH patients.**

**(A)** Differences in relative immune cell frequency across vascular regions in donor controls (Con) and pulmonary arterial hypertension (PAH) patients. IM, intramural; PV, perivascular; NV, non-vessel. **(B)** Immune cells distribution in PAH patients across vascular regions. **(C)** Schematics detailing how the overall vessel score was calculated for each patient are provided. The median of the pulmonary artery (PA) scores was calculated for each patient. This approach yielded a single value that reflects the central tendency of the vessel condition across different PAs, thus simplifying the assessment while accounting for intra-patient variability. **(D)** Heatmap displaying the correlation between immune cells across vascular regions (heatmap rows) and patient vessel scores (heatmap columns). The heatmap encompasses total cells (regardless of vascular region), as well as Intramural (IM), Perivascular (PV), and Non-Vessel (NV) regions. The p-values (white asterix) are reported. Spearman's rank correlation ($r_S$) was employed for the correlations, with a 95% confidence interval and two-tailed test. The p-values are reported as follows: *$p < 0.05$, **$p < 0.01$, ***$p < 0.001$, ****$p < 0.0001$. IM, intramural; VP, perivascular; NV, non-vessel; DC, dendritic cell; Mono, monocyte; Macro, macrophage; Neutro, neutrophil; Treg, regulatory T cell; NK, natural killer (NK) cell.

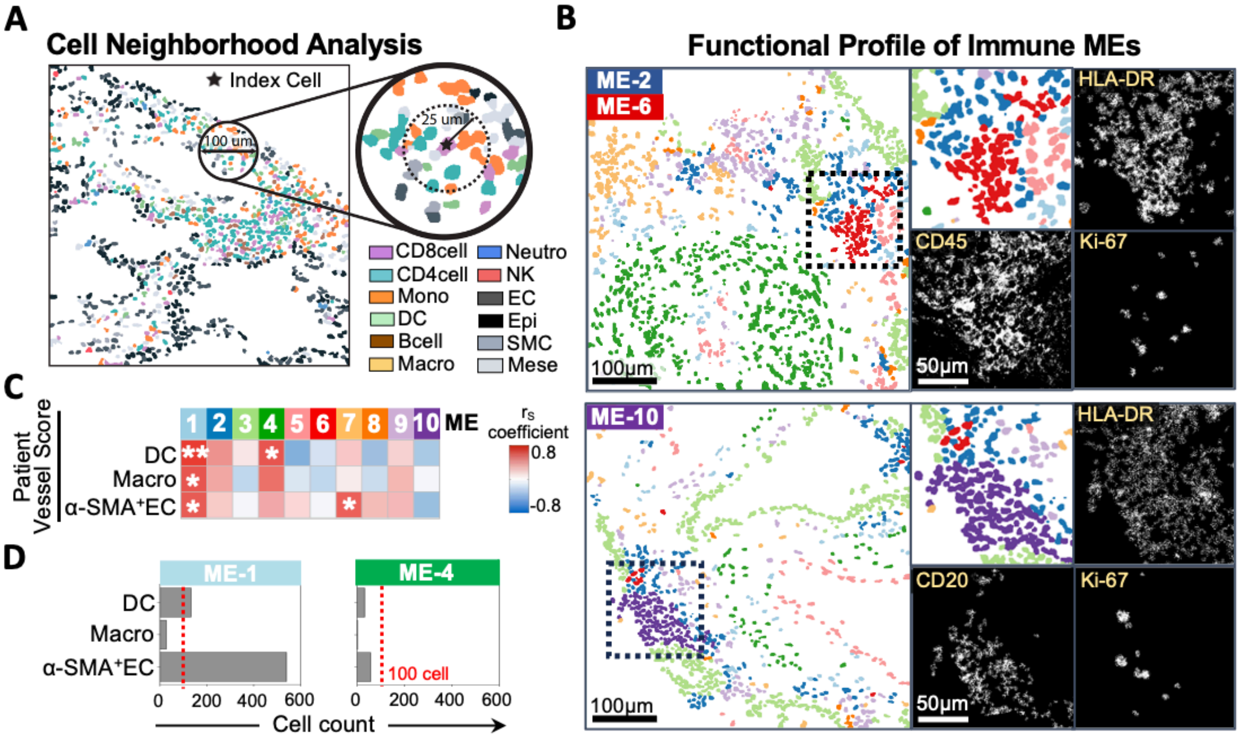


**Supplementary Figure E3. Mapping immune cell phenotypes and tissue microenvironments in pulmonary arteries.**

**(A)** Cell phenotypes are superimposed onto the segmentation mask of a representative pulmonary arterial section, creating a Cell Phenotype Map that illustrates the process of identifying tissue microenvironments (MEs). These MEs were determined based on the phenotypic traits and spatial coordinates of individual cells. This data was utilized to compute the occurrence rate of cell phenotypes located within approximately 25 µm of the centroid (index cell) of a specific cell.

**(B)** Functional profiles of immune-enriched MEs with a proliferative (Ki-67) and activated (HLA-DR) profile. The top panel displays ME-2 and ME-6, immune-enriched MEs surrounding the PAs. The bottom panel displays ME-10, a B cell-enriched ME. Images scale bars = 100 µm; zoomed inset scale bars = 50 µm. **(C)** Heatmap depicting the correlation between the frequency of DCs, macrophages and α-SMA+ endothelial cells in each tissue microenvironment (ME, heatmap columns) and patient vessel scores (heatmap rows). The p-values (white asterix) are reported.

Spearman's rank correlation ($r_S$) was employed for the correlations, with a 95% confidence interval and two-tailed test. **(D)** The absolute abundance of DCs, macrophages and α-SMA+ endothelial cells in ME-1 (left) and ME-4 (right). A dotted red line represents the threshold of 100 cells. The p-values are reported as follows: *$p < 0.05$, **$p < 0.01$, ***$p < 0.001$, ****$p < 0.0001$. DC, dendritic cell; Mono, monocyte; Macro, macrophage; Neutro, neutrophil; NK, natural killer (NK) cell; SMC, smooth muscle cell and fibroblast; Epi, epithelial cell; EC, endothelial cell; Mese, mesenchymal cell.

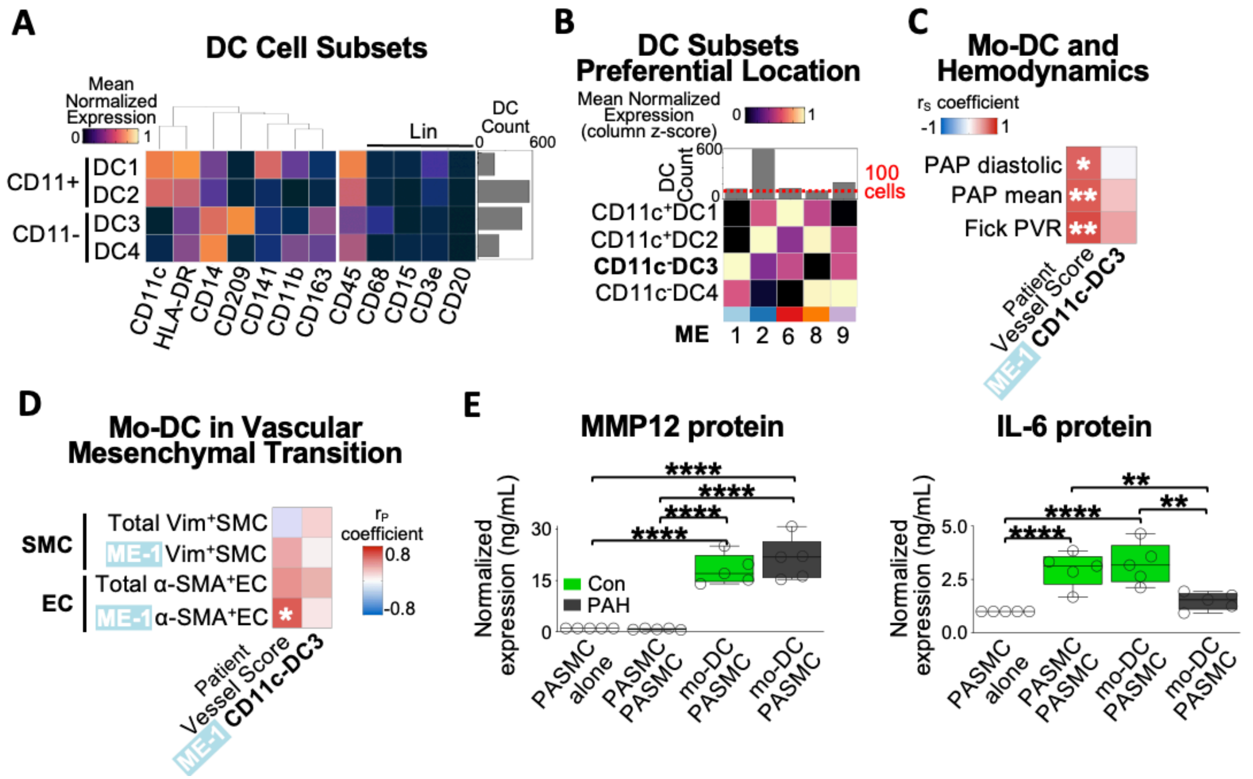


**Supplementary Figure E4. DC subsets and correlation analysis in PAH.**

**(A)** Dendritic cell assignments based on mean normalized expression of phenotypic markers (heatmap rows) hierarchically clustered (Euclidean distance, average linkage). The absolute abundance of each cell type is displayed (right). Lineage (Lin) markers were employed to differentiate dendritic cells (DCs) more accurately from T cells (CD3e), B cells (CD20), neutrophils (CD15), and macrophages (CD68). **(B)** Heatmap showing the z-scored mean normalized frequency of DC subsets across tissue microenvironments (MEs) in the heatmap rows. The absolute cell abundance within each ME is shown at the top. Only MEs with more than 100 DCs were included in the heatmap. A dotted red line represents the threshold of 100 cells. The CD11c-DC3s (mo-DC-like cells), highlighted in bold, were the predominant DC subtype in ME-1. **(C)** Heatmap depicting the correlation between hemodynamics (heatmap rows), and patient vessel scores or CD11c-DC3s within ME-1 (heatmap columns). PAP, pulmonary arterial pressure; PVR, pulmonary vascular resistance. The p-values (white asterix) are reported. Spearman's rank correlation ($r_S$) was employed for the correlations, with a 95% confidence interval and two-tailed test. **(D)** Heatmap depicting the correlation between the frequency of different phenotypes of SMCs and ECs found within the entire image 'Total' or within ME-1 (heatmap rows), and patient vessel scores or CD11c-DC3s within ME-1 (heatmap columns). The p-values (white asterix) are reported. Pearson correlation ($r_P$) was employed for the correlations, with a 95% confidence interval and two-tailed test. **(E)** Differences among culture conditions, involving monocyte-derived DCs (mo-DCs) and pulmonary arterial smooth muscle cells (PASMCs), for secreted matrix metalloproteinase-12 (MMP12) and IL-6 levels measured in cultured media. Five technical replicates are displayed per condition. Before evaluating co-culture differences, data was normalized to 'PASMC alone' control. Co-culture significance was assessed via a mixed-effects model, accounting for repeated measures as a random effect, followed by Tukey-adjusted pairwise comparisons. Significance levels are denoted as follows: *$p < 0.05$, **$p < 0.01$, ***$p < 0.001$, ****$p < 0.0001$. Con, donor control. DC, dendritic cell; Mono, monocyte; Macro, macrophage; Neutro, neutrophil; NK, natural killer (NK) cell.

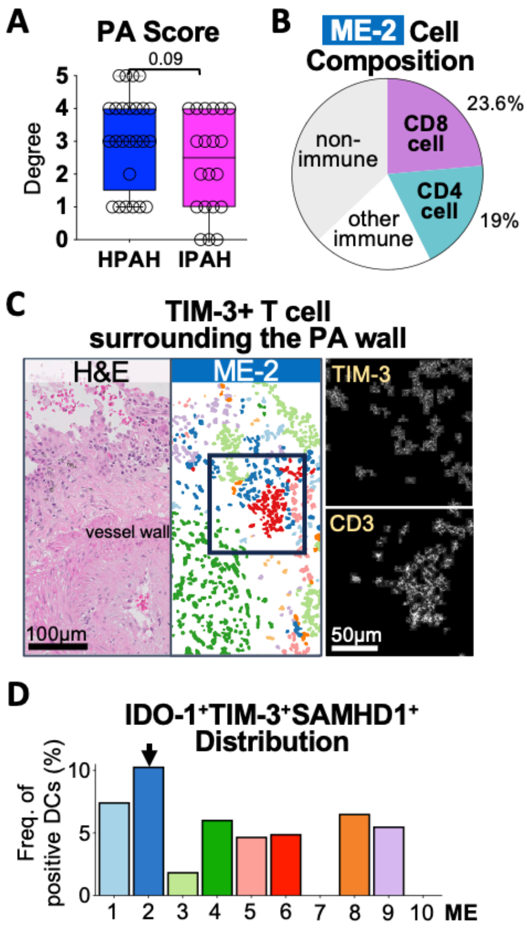


**Supplementary Figure E5. PAH subtypes and immune cell distribution.**

**(A)** Differences in pulmonary artery (PA) scores between PAH subtypes. **(B)** The cell composition of ME-2 is reported and highlights T cells, the most prevalent cell type. **(C)** Image set displaying TIM-3+ cells surrounding the pulmonary artery (PA) wall. On the left, half of the image displays the generated MEs, while the other half displays the hematoxylin and eosin (H&E) image to provide context within the PA tree. The zoomed inset highlights the perivascular location of ME-2. On the right, corresponding TIM-3 and CD3 images of the zoomed inset are shown. Images scale bars = 100 µm; zoomed inset scale bars = 50 µm. **(D)** The relative frequency of IDO-1+TIM-3+SAMHD1+ dendritic cells (DCs) across microenvironments (MEs). The accumulation of IDO-1+TIM-3+SAMHD1+ DCs in ME-2 is indicated with a black arrow. The p-values are reported as follows: *p < 0.05, **p < 0.01, ***p < 0.001, ****p < 0.0001.

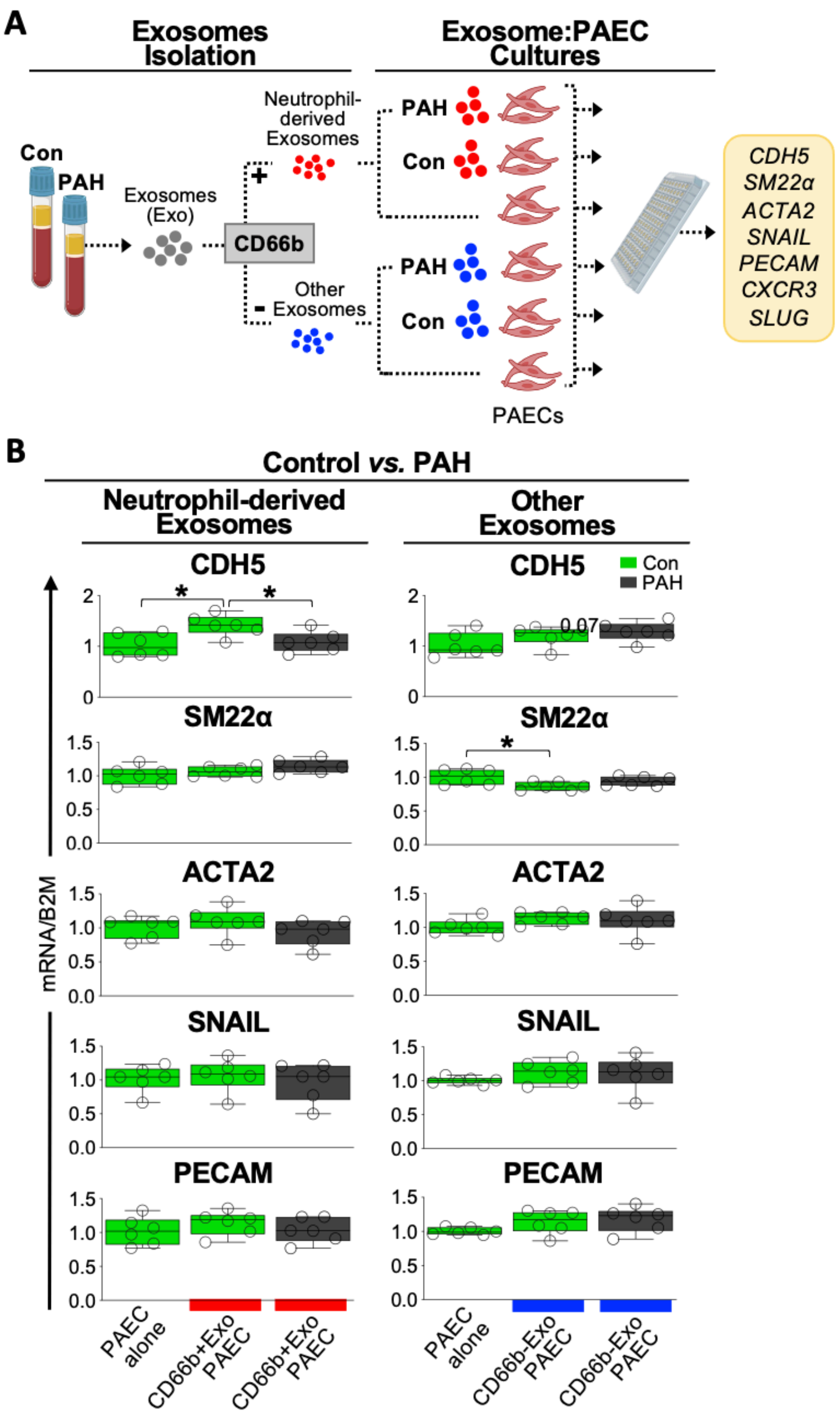


**Supplementary Figure E6. Neutrophil-derived exosome interaction with PAECs and gene expression analysis.**

**(A)** Schematic illustration depicting the workflow implemented to obtain and culture exosomes (Exo) derived from neutrophils (CD66b+Exo) as well as exosomes from other sources (CD66b-

Exo) with pulmonary arterial endothelial cells (PAECs) from donor controls (Con) and PAH patients. Cadherin-5 (CDH5), Smooth Muscle Protein 22-Alpha (SM22α), Actin Alpha Cardiac Muscle 2 (ACTA2), Snail Family Transcriptional Repressor 1 (SNAIL or SNAI1), Platelet Endothelial Cell Adhesion Molecule (PECAM), C-X-C chemokine receptor type 3 (CXCR3), and Snail Family Transcriptional Repressor 2 (SLUG or SNAI2) gene expression were measured in pulmonary arterial endothelial cells (PAECs). **(B)** Differences among culture conditions, involving exosomes (Exo) and pulmonary arterial endothelial cells (PAECs), were assessed for CDH5, SM22α, ACTA2, PECAM, and SNAIL gene in pulmonary arterial endothelial cells (PAECs). CXCR3 and SLUG are not displayed given their high ct. Six technical replicates are displayed per condition. Con, donor controls. Before evaluating co-culture differences, data was normalized to 'PAEC alone' control. Co-culture significance was assessed via a mixed-effects model, accounting for repeated measures as a random effect, followed by Tukey-adjusted pairwise comparisons. Significance levels are denoted as follows: $^*p < 0.05$, $^{**}p < 0.01$, $^{***}p < 0.001$, $^{****}p < 0.0001$.

### Table E1

| Target | Clone | Channel | Provider | Catalogue |
|---|---|---|---|---|
| **Antibody cocktail 1** | | | | |
| HLA Class I | EMR8-5 | 141Pr | AbCam | ab70328 |
| CD4 | EPR6855 | 143Nd | AbCam | ab181724 |
| CD14 | D7A2T | 144Nd | CST | 56082BF |
| FoxP3 | 236A/E7 | 146Nd | BD Biosciences | 624084 |
| PD-1 | D4W2J | 147Sm | CST | 86163BF |
| CD31 | EP3095 | 148Nd | AbCam | ab216459 |
| PD-L1 biotinylated | E1L3N | 149Sm | CST | 15118S |
| Granzyme B | D6E9W | 150Nd | IONPath | 715002 |
| Ki-67 | 8D5 | 151Eu | CST | 9449BF |
| CD209 | DCN46 | 152Sm | BD Biosciences | 624084 |
| CD45RO | UCHL1 | 153Eu | Biolegend | 304202 |
| SAMHD1 | Polyclonal | 154Sm | AbCam | ab67820 |
| iNOS | SP126 | 155Gd | Spring Bioscience | M4264 |
| CD68 | D4B9C | 156Gd | Biolegend | 304202 |
| CD141 | EPR4051 | 157Gd | AbCam | ab222292 |
| CD8 | C8/144B | 158Gd | Cell Marque | 108M-9 |
| CD3e | D7A6E | 159Tb | CST | 85061BF |
| CD15 | W6D3 | 160Gd | BD Biosciences | 557895 |
| CD11c | EP1347Y | 161Dy | AbCam | ab216655 |
| TIM-3 | EPR22241 | 162Dy | AbCam | ab242080 |
| CD163 | D6U1J | 163Dy | CST | 93498BF |
| CD20 | L26 | 164Dy | Cell Marque | 120M-8 |
| CD16 | D1N9L | 165Ho | CST | 24326BF |
| IFN-γ | IFNG466 | 166Er | AbCam | ab218890 |
| HLA-DR | EPR3692 | 167Er | AbCam | ab215985 |
| CD11b | EP1345Y | 168Er | AbCam | ab187537 |
| CD45 | D9M8I | 169Tm | CST | 13917BF |
| IDO-1 | SP260 | 170Er | Spring Bioscience | M5604.C |
| Pan-cytokeratin | AE1/AE3 | 171Yb | ThermoFisher | MS-343-PABX |
| b-Catenin | D10A8 | 172Yb | CST | 8480BF |
| Na+/K+ ATPase | EP1845Y | 175Yb | AbCam | ab167390 |
| MPO | polyclonal | 176Yb | R&D System | AF3667 |
| **Antibody cocktail 2** | | | | |
| Histone H3 | D1H2 | 89Y | CST | 4499BF |
| Vimentin | D21H3 | 113In | CST | 5741BF |
| Smooth Muscle Actin | D4K9N | 115In | CST | 19245BF |
| Biotin | 1D4-C5 | 149Sm | Biolegend | 409002 |

**Supplementary Table E1. Antibody panel.**

List of antibodies, clones, metal isotypes, vendors and catalogue number used to make up the polychromatic panel. Antibody cocktail 2 was used for second day staining. Y, Yttrium; In, Indium; Pr, Praseodymium; Nd, Neodymium, Sm, Samarium; Eu, Europium; Gd, Gadolinium; Tb, Terbium; Dy, Dysprosium; Er, Erbium; Tm, Thulium; Yb, Ytterbium.

### Table E2

| Predictor | Estimate | SE Coef | z value | OR | CI | | Pr (>\|z\|) |
|---|---|---|---|---|---|---|---|
| *PA microenvironments* | | | | | | | |
| ME-1 (%) | 0.212 | 0.053 | 4.003 | 1.237 | 0.115 | 0.324 | 6.25e-05 **** |
| ME-2 (%) | 0.007 | 0.030 | 0.235 | 1.007 | -0.053 | 0.068 | 0.814 |
| ME-3 (%) | 0.001 | 0.027 | 0.034 | 1.000 | -0.053 | 0.056 | 0.973 |
| ME-4 (%) | 0.056 | 0.030 | 1.869 | 1.057 | -0.001 | 0.117 | 0.061 |
| ME-5 (%) | -0.060 | 0.030 | -1.946 | 0.941 | -0.122 | 0.001 | 0.051 |
| ME-6 (%) | -0.024 | 0.021 | -1.166 | 0.975 | -0.071 | 0.016 | 0.244 |
| ME-7 (%) | -0.011 | 0.023 | -0.466 | 0.988 | -0.057 | 0.038 | 0.641 |
| ME-8 (%) | -0.006 | 0.027 | -0.228 | 0.993 | -0.061 | 0.046 | 0.819 |
| ME-9 (%) | -0.010 | 0.030 | -0.337 | 0.989 | -0.071 | 0.050 | 0.736 |
| ME-10 (%) | 0.042 | 0.045 | -0.943 | 0.958 | -0.134 | 0.057 | 0.346 |

**Supplementary Table E2. Tissue microenvironments as predictors of PA score.**

Independent contribution of the ten-tissue microenvironment (ME) to variations in the PA score. A summary of the outcomes of an ordinal regression analysis is reported, where each ME, defined as the frequency of cells within a specific microenvironment of the pulmonary artery (PA), relative to the total number of cells in the PA, serves as an independent predictor of the PA score. The coefficient estimates (Estimate) for each microenvironment (predictor variable) indicate the direction and strength of their association with the PA score. The standard error coefficient (SE Coef) provides information about the precision of the estimated coefficients. The z-values offer insights into the statistical significance, direction, and magnitude of the effect of each ME on the PA score. The odds ratios (OR) along with their 95% confidence intervals (CI), quantify the change in the odds of a higher PA score associated with a one-unit change in the predictor variable. The p-value associated with the value (Pr (>|z|) indicates the statistical significance of each predictor variable, where *p < 0.05, **p < 0.01, ***p < 0.001, ****p < 0.0001.

### Table E3

| Gene | Mean (SD) Condition A | Condition B | Condition C | Condition D | Post Hoc Test | Estimate | SE | df | t ratio | Adjusted P-value |
|---|---|---|---|---|---|---|---|---|---|---|
| ACTA2 | 1.024 (0.245) | 0.950 (0.326) | 1.334 (0.272) | 1.279 (0.299) | A-B | 0.075 | 0.177 | 12 | 0.422 | 0.974 |
| | | | | | A-C | -0.310 | 0.177 | 12 | -1.753 | 0.341 |
| | | | | | A-D | -0.255 | 0.177 | 12 | -1.440 | 0.500 |
| | | | | | B-C | -0.385 | 0.177 | 12 | -2.175 | 0.185 |
| | | | | | B-D | -0.329 | 0.177 | 12 | -1.862 | 0.294 |
| | | | | | C-D | 0.055 | 0.177 | 12 | 0.312 | 0.989 |
| CD14 # | 1.108 (0.411) | 0.747 (0.311) | 1.292 (0.805) | 1.057 (0.451) | A-B | 0.361 | 0.334 | 12 | 1.080 | 0.708 |
| | | | | | A-C | -0.184 | 0.334 | 12 | -0.551 | 0.945 |
| | | | | | A-D | 0.051 | 0.334 | 12 | 0.151 | 0.999 |
| | | | | | B-C | -0.545 | 0.334 | 12 | -1.631 | 0.399 |
| | | | | | B-D | -0.310 | 0.334 | 12 | -0.929 | 0.790 |
| | | | | | C-D | 0.235 | 0.334 | 12 | 0.702 | 0.894 |
| Fibronectin | 1.041 (0.256) | 0.953 (0.137) | 1.193 (0.193) | 1.217 (0.204) | A-B | 0.088 | 0.123 | 12 | 0.721 | 0.887 |
| | | | | | A-C | -0.152 | 0.123 | 12 | -1.241 | 0.615 |
| | | | | | A-D | -0.176 | 0.123 | 12 | -1.436 | 0.502 |
| | | | | | B-C | -0.241 | 0.123 | 12 | -1.962 | 0.255 |
| | | | | | B-D | -0.265 | 0.123 | 12 | -2.157 | 0.191 |
| | | | | | C-D | -0.024 | 0.123 | 12 | -0.196 | 0.997 |
| MMP12 | 1.080 (0.288) | 1.768 (0.438) | 1.250 (0.302) | 0.756 (0.190) | A-B | -0.688 | 0.169 | 12 | -4.074 | 0.007 ** |
| | | | | | A-C | -0.169 | 0.169 | 12 | -1.004 | 0.750 |
| | | | | | A-D | 0.324 | 0.169 | 12 | 1.919 | 0.271 |
| | | | | | B-C | 0.519 | 0.169 | 12 | 3.070 | 0.042 * |
| | | | | | B-D | 1.012 | 0.169 | 12 | 5.993 | 0.0003 *** |
| | | | | | C-D | 0.494 | 0.169 | 12 | 2.923 | 0.054 |
| SM22α | 1.012 (0.168) | 0.960 (0.061) | 1.009 (0.133) | 0.960 (0.022) | A-B | 0.053 | 0.071 | 12 | 0.742 | 0.878 |
| | | | | | A-C | 0.004 | 0.071 | 12 | 0.056 | 0.999 |
| | | | | | A-D | 0.053 | 0.071 | 12 | 0.744 | 0.878 |
| | | | | | B-C | -0.049 | 0.071 | 12 | -0.686 | 0.900 |
| | | | | | B-D | 0.0001 | 0.071 | 12 | 0.002 | 1 |
| | | | | | C-D | 0.098 | 0.071 | 12 | 0.688 | 0.900 |
| Vimentin | 1.015 (0.185) | 0.934 (0.219) | 1.221 (0.275) | 1.175 (0.165) | A-B | 0.082 | 0.121 | 12 | 0.677 | 0.904 |
| | | | | | A-C | -0.206 | 0.121 | 12 | -1.703 | 0.364 |
| | | | | | A-D | -0.159 | 0.121 | 12 | -1.319 | 0.569 |
| | | | | | B-C | -0.287 | 0.121 | 12 | -2.380 | 0.134 |
| | | | | | B-D | -0.241 | 0.121 | 12 | -1.996 | 0.243 |
| | | | | | C-D | 0.046 | 0.121 | 12 | 0.384 | 0.980 |
| CXCR3 # | 1.122 (0.660) | 1.044 (0.340) | 1.082 (0.2233) | 0.8316 (0.1389) | A-B | 0.078 | 0.249 | 12 | 0.313 | 0.989 |
| | | | | | A-C | 0.040 | 0.249 | 12 | 0.160 | 0.998 |
| | | | | | A-D | 0.290 | 0.249 | 12 | 1.167 | 0.658 |
| | | | | | B-C | -0.038 | 0.249 | 12 | -0.153 | 0.999 |
| | | | | | B-D | 0.213 | 0.249 | 12 | 0.854 | 0.828 |
| | | | | | C-D | 0.251 | 0.249 | 12 | 1.007 | 0.748 |
| MMP9 | Not detected | | | | | | | | | |
| INOS | Not detected | | | | | | | | | |

**Supplementary Table E3. Downregulation of MMP12 gene in monocyte-derived DCs-stimulated pulmonary arterial smooth muscle cells.**

Gene expression changes in pulmonary arterial smooth muscle cells (PASMCs) in response to co-cultures with donor vs. PAH monocyte-derived dendritic cells (mo-DCs). A list of transcripts associated with the shift in PASMC phenotype was assessed using RT-qPCR across four culture conditions: Condition A: PASMCs cultured alone. Condition B: PASMCs co-cultured with other PASMCs. Condition C: PASMCs co-cultured with donor mo-DCs. Condition D: PASMCs co-cultured with PAH mo-DCs. The symbol '#', highlights genes with higher Ct values due to low levels in the sample.

The mean and standard deviation (SD) of five technical replicates across genes and culture conditions are reported. Prior to evaluating co-culture differences, data were normalized to the PASMC alone control. The significance of co-culture effects was assessed using a mixed-effects model, accounting for repeated measures as a random effect. Pairwise comparisons between the estimated marginal means (EMMs) were conducted, and Tukey adjustment was applied to control the family-wise error rate. This model aims to detect differences in transcript expression between the various co-culture conditions while considering the variability between technical replicates within each condition. The results are presented as estimate coefficient (estimate), standard error (SE), degrees of freedom (df), t-ratio, and P-values. The P-value (Adjusted P-value) was obtained after correcting for multiple comparisons using the Tukey adjustment and is represented as follows: <0.0001 '****' <0.001 '***' <0.01 '**' <0.05 '*'.

### Table E4

| Signaling Molecule | Mean (SD) Condition A | Condition B | Condition C | Condition D | Post Hoc Test | Estimate | SE | df | t ratio | Adjusted P value |
|---|---|---|---|---|---|---|---|---|---|---|
| TNF-α | 0.000 (0.000) | 0.000 (0.000) | 21.130 (4.537) | 4.786 (2.578) | A-B | 0.000 | 1.456 | 12 | 0.000 | 1.000 |
| | | | | | A-C | -21.129 | 1.456 | 12 | -14.513 | 0.000 **** |
| | | | | | A-D | -4.786 | 1.456 | 12 | -3.288 | 0.029 * |
| | | | | | B-C | -21.129 | 1.456 | 12 | -14.513 | 0.000 **** |
| | | | | | B-D | -4.786 | 1.456 | 12 | -3.288 | 0.029 * |
| | | | | | C-D | 16.343 | 1.456 | 12 | 11.225 | 0.000 **** |
| IL-6 | 1273 (305.000) | 3584 (423.500) | 3971 (882.200) | 1800 (219.700) | A-B | -1.965 | 0.35 | 12 | -5.612 | <0.001 **** |
| | | | | | A-C | 2.227 | 0.35 | 12 | 6.362 | <0.001 **** |
| | | | | | A-D | 0.484 | 0.35 | 12 | 1.382 | 0.533 |
| | | | | | B-C | 0.263 | 0.35 | 12 | 0.750 | 0.875 |
| | | | | | B-D | -1.481 | 0.35 | 12 | -4.230 | 0.006 ** |
| | | | | | C-D | 1.743 | 0.35 | 12 | 4.980 | 0.002 ** |
| MMP12 | 1.122 (0.660) | 1.044 (0.340) | 1.082 (0.2233 | 0.8316 0.1389 | A-B | -0.636 | 2.181 | 11 | -0.291 | 0.991 |
| | | | | | A-C | -18.072 | 2.181 | 11 | -8.285 | 0.000**** |
| | | | | | A-D | -22.547 | 2.353 | 11 | -9.583 | 0.000**** |
| | | | | | B-C | -17.436 | 2.181 | 11 | -7.994 | 0.000**** |
| | | | | | B-D | -21.911 | 2.353 | 11 | -9.312 | 0.000**** |
| | | | | | C-D | -4.475 | 2.353 | 11 | -1.902 | 0.279 |
| IFN-α | Not detected | | | | | | | | | |
| IFN-β | Not detected | | | | | | | | | |
| IFN-γ | Not detected | | | | | | | | | |
| IL-12 | Not detected | | | | | | | | | |
| IL-1β | Not detected | | | | | | | | | |
| IP-10 | Not detected | | | | | | | | | |

**Supplementary Table E4. Differential signaling molecules in donor and PAH monocyte-derived DCs co-cultured with pulmonary arterial smooth muscle cells.**

Signaling molecule changes in culture media between donor monocyte-derived dendritic cells (mo-DCs):PASMC and PAH mo-DCs:PASMC co-cultures. A list of signaling molecules, including cytokines and a chemokine receptor, known to impact PASMCs and derived from mo-DCs, was assessed using ELISA across four culture conditions: Condition A: PASMCs cultured alone. Condition B: PASMCs co-cultured with other PASMCs. Condition C: PASMCs co-cultured with donor mo-DCs. Condition D: PASMCs co-cultured with PAH mo-DCs.

The mean and standard deviation (SD) of five technical replicates across signaling molecules and culture conditions are reported. Prior to evaluating co-culture differences, data were normalized to the PASMC alone control. The significance of co-culture effects was assessed using a mixed-effects model, accounting for repeated measures as a random effect. Pairwise comparisons between the estimated marginal means (EMMs) were conducted, and Tukey adjustment was applied to control the family-wise error rate. This model aims to detect differences in signaling molecule levels between the various co-culture conditions while considering the variability between technical replicates within each condition. The results are presented as estimate coefficient (estimate), standard error (SE), degrees of freedom (df), t-ratio, and P-values. The P-value (Adjusted P-value) was obtained after correcting for multiple comparisons using the Tukey adjustment and is represented as follows: <0.0001 '****' <0.001 '***' <0.01 '**' <0.05 '*'.

### Table E5

| Gene | Mean (SD) Condition A | Condition B | Condition C | Condition D | Post Hoc Test | Estimate | SE | df | t ratio | Adjusted P-value |
|---|---|---|---|---|---|---|---|---|---|---|
| PECAM | 1.007 (0.124) | 0.968 (0.024) | 0.655 (0.174) | 0.697 (0.166) | A-B | 0.039 | 0.098 | 9.551 | 0.401 | 0.977 |
| | | | | | A-C | 0.352 | 0.098 | 9.551 | 3.603 | 0.022 * |
| | | | | | A-D | 0.310 | 0.093 | 10.114 | 3.325 | 0.032 * |
| | | | | | B-C | 0.313 | 0.098 | 9.551 | 3.203 | 0.042 * |
| | | | | | B-D | 0.271 | 0.093 | 10.114 | 2.905 | 0.063 |
| | | | | | C-D | -0.042 | 0.093 | 10.114 | -0.450 | 0.968 |
| SM22α | 1.001 (0.055) | 0.783 (0.013) | 0.593 (0.105) | 0.705 (0.213) | A-B | 0.219 | 0.091 | 9.511 | 2.398 | 0.143 |
| | | | | | A-C | 0.409 | 0.091 | 9.511 | 4.481 | 0.006 ** |
| | | | | | A-D | 0.298 | 0.087 | 10.109 | 3.406 | 0.028 * |
| | | | | | B-C | 0.190 | 0.091 | 9.511 | 2.082 | 0.226 |
| | | | | | B-D | 0.079 | 0.087 | 10.109 | 0.901 | 0.805 |
| | | | | | C-D | -0.111 | 0.087 | 10.109 | -1.274 | 0.598 |
| CDH5 | 1.025 (0.245) | 1.139 (0.158) | 0.671 (0.107) | 0.620 (0.114) | A-B | -0.115 | 0.099 | 9.299 | -1.164 | 0.662 |
| | | | | | A-C | 0.354 | 0.099 | 9.299 | 3.586 | 0.024 * |
| | | | | | A-D | 0.408 | 0.096 | 10.012 | 4.239 | 0.008 ** |
| | | | | | B-C | 0.469 | 0.099 | 9.299 | 4.750 | 0.004 ** |
| | | | | | B-D | 0.522 | 0.096 | 10.012 | 5.433 | 0.001 ** |
| | | | | | C-D | 0.054 | 0.096 | 10.012 | 0.560 | 0.942 |
| CXCR3 # | 1.030 (0.270) | 0.814 (0.144) | 0.858 (0.309) | 0.878 (0.324) | A-B | 0.216 | 0.195 | 9.551 | 1.109 | 0.693 |
| | | | | | A-C | 0.173 | 0.195 | 9.551 | 0.886 | 0.812 |
| | | | | | A-D | 0.153 | 0.186 | 10.114 | 0.821 | 0.843 |
| | | | | | B-C | -0.043 | 0.195 | 9.551 | -0.222 | 0.996 |
| | | | | | B-D | -0.063 | 0.186 | 10.114 | -0.340 | 0.986 |
| | | | | | C-D | -0.020 | 0.186 | 10.114 | -0.107 | 1.000 |
| SLUG # | 1.014 (0.110) | 0.601 (0.082) | 0.684 (0.194) | 0.762 (0.273) | A-B | 0.413 | 0.131 | 9.495 | 3.163 | 0.045 |
| | | | | | A-C | 0.330 | 0.131 | 9.495 | 2.526 | 0.119 |
| | | | | | A-D | 0.252 | 0.125 | 10.106 | 2.013 | 0.246 |
| | | | | | B-C | -0.083 | 0.131 | 9.495 | -0.637 | 0.917 |
| | | | | | B-D | -0.161 | 0.125 | 10.106 | -1.287 | 0.590 |
| | | | | | C-D | -0.078 | 0.125 | 10.106 | -0.622 | 0.923 |
| SNAIL | 1.009 (0.101) | 0.849 (0.067) | 0.780 (0.253) | 0.912 (0.280) | A-B | 0.160 | 0.140 | 9.470 | 1.146 | 0.672 |
| | | | | | A-C | 0.229 | 0.140 | 9.470 | 1.632 | 0.407 |
| | | | | | A-D | 0.099 | 0.135 | 10.100 | 0.734 | 0.881 |
| | | | | | B-C | 0.068 | 0.140 | 9.470 | 0.486 | 0.960 |
| | | | | | B-D | -0.062 | 0.135 | 10.100 | -0.459 | 0.966 |
| | | | | | C-D | -0.130 | 0.135 | 10.100 | -0.966 | 0.771 |
| ACTA2 | 1.007 (0.127) | 0.898 (0.095) | 0.771 (0.212) | 0.921 (0.230) | A-B | 0.110 | 0.128 | 9.551 | 0.860 | 0.825 |
| | | | | | A-C | 0.236 | 0.128 | 9.551 | 1.852 | 0.309 |
| | | | | | A-D | 0.087 | 0.122 | 10.114 | 0.713 | 0.890 |
| | | | | | B-C | 0.126 | 0.128 | 9.551 | 0.992 | 0.758 |
| | | | | | B-D | -0.023 | 0.122 | 10.114 | -0.188 | 0.997 |
| | | | | | C-D | -0.149 | 0.122 | 10.114 | -1.227 | 0.625 |
| INOS | Not detected | | | | | | | | | |

**Supplementary Table E5. Downregulation of PECAM and CDH5 gene in monocyte-derived DCs-stimulated pulmonary arterial endothelial cells.**

Gene expression changes in pulmonary arterial endothelial cells (PAECs) in response to co-cultures with donor vs. PAH monocyte-derived dendritic cells (mo-DCs). A list of transcripts associated with the shift in PAECs phenotype was assessed using RT-qPCR across four culture conditions: Condition A: PAECs cultured alone. Condition B: PAECs co-cultured with other PAECs. Condition C: PAECs co-cultured with donor mo-DCs. Condition D: PAECs co-cultured with PAH mo-DCs. The symbol '#', highlights genes with higher Ct values due to low levels in the sample.

The mean and standard deviation (SD) of five technical replicates across genes and culture conditions are reported. Prior to evaluating co-culture differences, data were normalized to the PAECs alone control. The significance of co-culture effects was assessed using a mixed-effects model, accounting for repeated measures as a random effect. Pairwise comparisons between the estimated marginal means (EMMs) were conducted, and Tukey adjustment was applied to control the family-wise error rate. This model aims to detect differences in transcript expression between the various co-culture conditions while considering the variability between technical replicates within each condition. The results are presented as estimate coefficient (estimate), standard error (SE), degrees of freedom (df), t-ratio, and P-values. The P-value (Adjusted P-value) was obtained after correcting for multiple comparisons using the Tukey adjustment and is represented as follows: <0.0001 '****' <0.001 '***' <0.01 '**' <0.05 '*'.
